# Surface condensation of a pioneer transcription factor on DNA

**DOI:** 10.1101/2020.09.24.311712

**Authors:** Jose A. Morin, Sina Wittmann, Sandeep Choubey, Adam Klosin, Stefan Golfier, Anthony A. Hyman, Frank Jülicher, Stephan W. Grill

**Affiliations:** Max Planck Institute of Molecular Cell Biology and Genetics, Pfotenhauerstraße 108, Dresden, Germany; Biotechnologisches Zentrum, Technische Universität Dresden, Tatzberg 47/49, Dresden, Germany; Max Planck Institute for the Physics of Complex Systems, Nöthnitzer Straße 38, Dresden, Germany; Center for Systems Biology Dresden, Pfotenhauerstraße 108, Dresden, Germany; Cluster of Excellence Physics of Life, Technische Universität Dresden, Dresden, Germany

## Abstract

In the last decade, extensive studies on the properties of non-membrane-bound compartments in the cellular cytoplasm have shown that concepts in phase separation drawn from physical chemistry can describe their formation and behaviour^1–4^. Current evidence also suggests that phase separation plays a role in the organization inside the cell nucleus^5–8^. However, the influence and role of DNA on the physical chemistry of phase separation is not well understood. Here, we are interested in the role of interactions between phase separating proteins and the DNA surface. The interaction of liquid phases with surfaces has been extensively studied in soft matter physics, in the context of macroscopic surfaces and non-biological liquids^9–11^. The conditions in the nucleus are different from those studied in conventional soft matter physics because DNA with a diameter of about 2 nm^12^ provides a microscopic surface, and liquid-like phases are complex mixtures of proteins subject to a myriad of biochemical modifications^13^. Transcriptional condensates, which are thought to serve as regulatory hubs in gene expression^14–21^, provide an accessible system to investigate the physics of condensates that emerge from DNA-protein and protein-protein interactions. These condensates are typically small^22^, and the mechanisms that determine their size are unknown. Whether they can be understood as phase separated compartments has been subject to debate^23–26^. Here, we use optical tweezers to directly observe the condensation of the pioneer transcription factor Klf4^27,28^ on DNA *in vitro*. We demonstrate that Klf4 forms microphases that are enabled by interaction with the DNA surface. This sets their typical size and allows them to form below the saturation concentration for liquid-liquid phase separation. We combine experiment with theory to show that these microphases can be understood as forming by surface condensation on DNA via a switch-like transition similar to prewetting. Polymer surface mediated condensation reconciles several observations that were previously thought to be at odds with the idea of phase separation as an organizing principle in the nucleus.

---

We focused on the human pioneer factor Klf4 (Krüppel-like factor 4), a driver of differentiation, cell growth and proliferation^29^ which is also one of the Yamanaka factors used for cell reprogramming into a pluripotent state^30^. Klf4 has a domain organization typical for transcription factors: an activation domain predicted to be disordered, and a structured DNA-binding region^31^. Human Klf4 with a C-terminal GFP tag purified from insect cells^32^ (Fig. 1a, Extended Data Fig. 1a) binds to DNA oligonucleotides in a sequence specific manner, as expected (Fig. 1b, Extended Data Fig. 2). Klf4 could form condensates in the absence of DNA above a concentration of ~1.2 μM in bulk solution (Fig. 1c top row, Extended Data Fig. 3a), which is above the estimated nuclear concentration of Klf4 (approximately 100 nM, Extended Data Fig. 1b). These condensates have behaviours of liquid-like drops (Fig. 1d, Extended Data Fig. 3b). Untagged Klf4 formed condensates at similar concentrations (Extended Data Fig. 3c, d and e). The addition of λ-DNA triggered the formation of foci below the saturation concentration (Fig. 1c bottom row), confirming previous observations that DNA can trigger foci formation of transcription factors^33^.

**Fig. 1.**
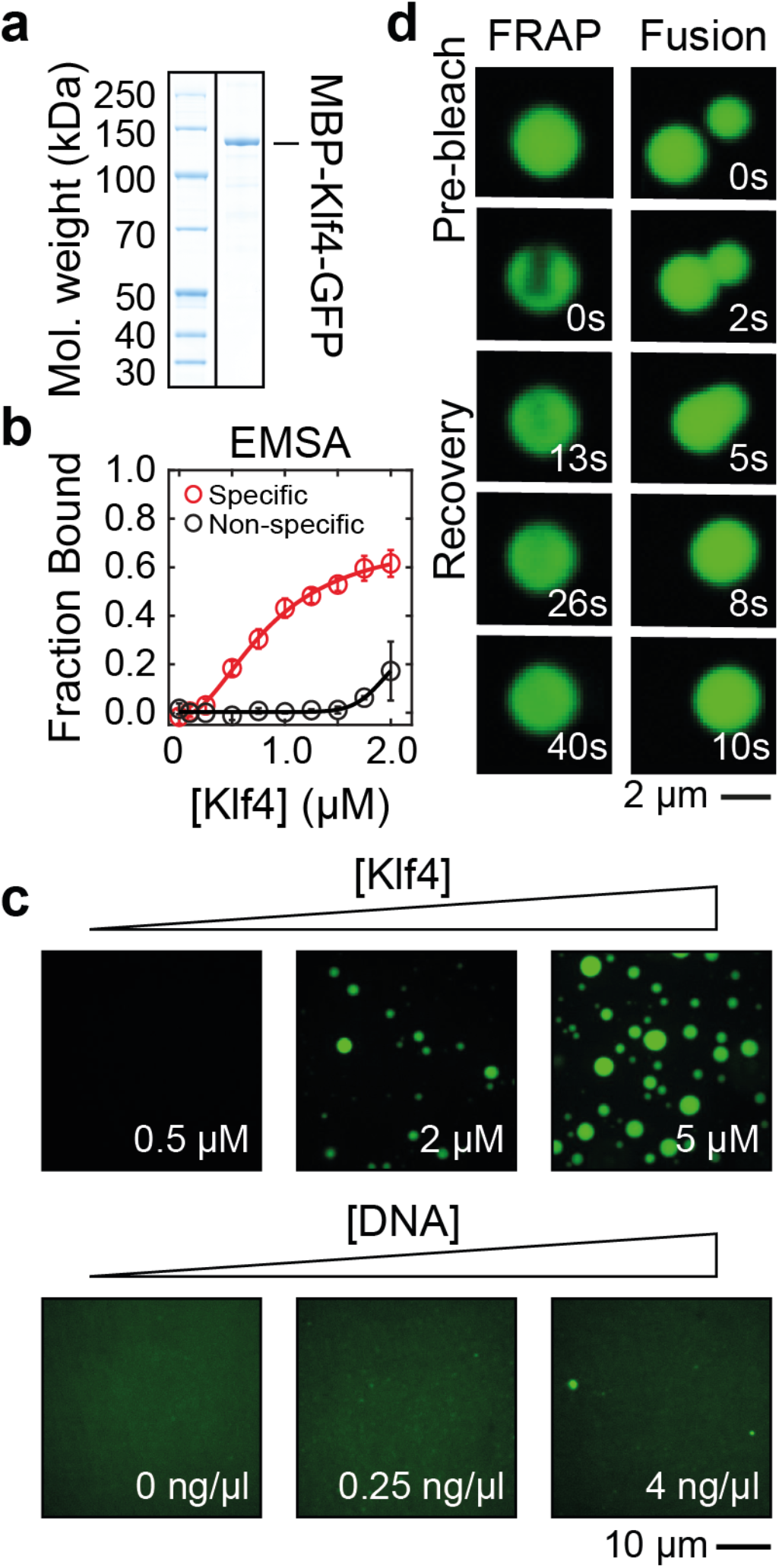
Klf4 forms phase separated condensates *in vitro*. **a**, SDS gel showing recombinantly expressed and purified MBP-Klf4-GFP. **b**, Test of the affinity of Klf4-GFP to short dsDNA oligonucleotides in the presence (red) or absence (black) of specific binding sites using electrophoretic mobility shift assays (EMSA; see also Extended Data Fig. 2). Error bars, standard deviation of three independent experiments. **c**, Top row, phase separation assay of Klf4-GFP reveals bulk droplet formation above a saturation concentration of ~1.2 μM (see Extended Data Fig. 3a). Bottom row, the addition of λ-DNA to 750 nM of Klf4-GFP triggers foci formation. **d**, Confocal microscopy images of Klf4-GFP droplets reveal liquid-like properties, as assessed by fluorescence recovery after photobleaching (FRAP; see also Extended Data Fig. 3b) and droplet fusion.

To further examine the behaviour of Klf4 on DNA, we used a dual trap optical tweezer to hold a linearized λ-DNA molecule stretched between two polystyrene beads (Fig. 2a)^34–37^. A combination of fluorescence readout and controlled microfluidics allowed us to detect interactions between Klf4 and DNA^38^. We observed many Klf4 foci on the DNA molecule (Fig. 2b and Supplementary Video 1) at a Klf4 concentration of 115 nM, which is similar to the estimated nuclear concentration (Extended Data Fig. 1b). These regions varied in the amount of Klf4 they contain, but they grew with time and reached a finite size with an average of approximately 800 molecules per cluster (Fig. 2d, Extended Data Fig. 4a-e). Furthermore, they can fuse and their position can fluctuate on DNA (Fig. 2c and Extended Data Fig. 6). Taken together, these observations suggest that Klf4 can form small condensate-like objects on DNA that grow to a finite size. We verified that these condensates form well below the saturation concentration of the bulk solution (Extended Data Fig. 3a and 7a). It is important to note that because condensates form on DNA well below the saturation concentration in the bulk, this excludes a scenario in which DNA serves as a nucleator for the formation of bulk phase droplets (Extended Data Fig. 7b). This is because in this scenario, bulk droplets could only grow if the solution remains above the saturation concentration (Extended Data Fig. 8)^33,39,40^.

**Fig. 2.**
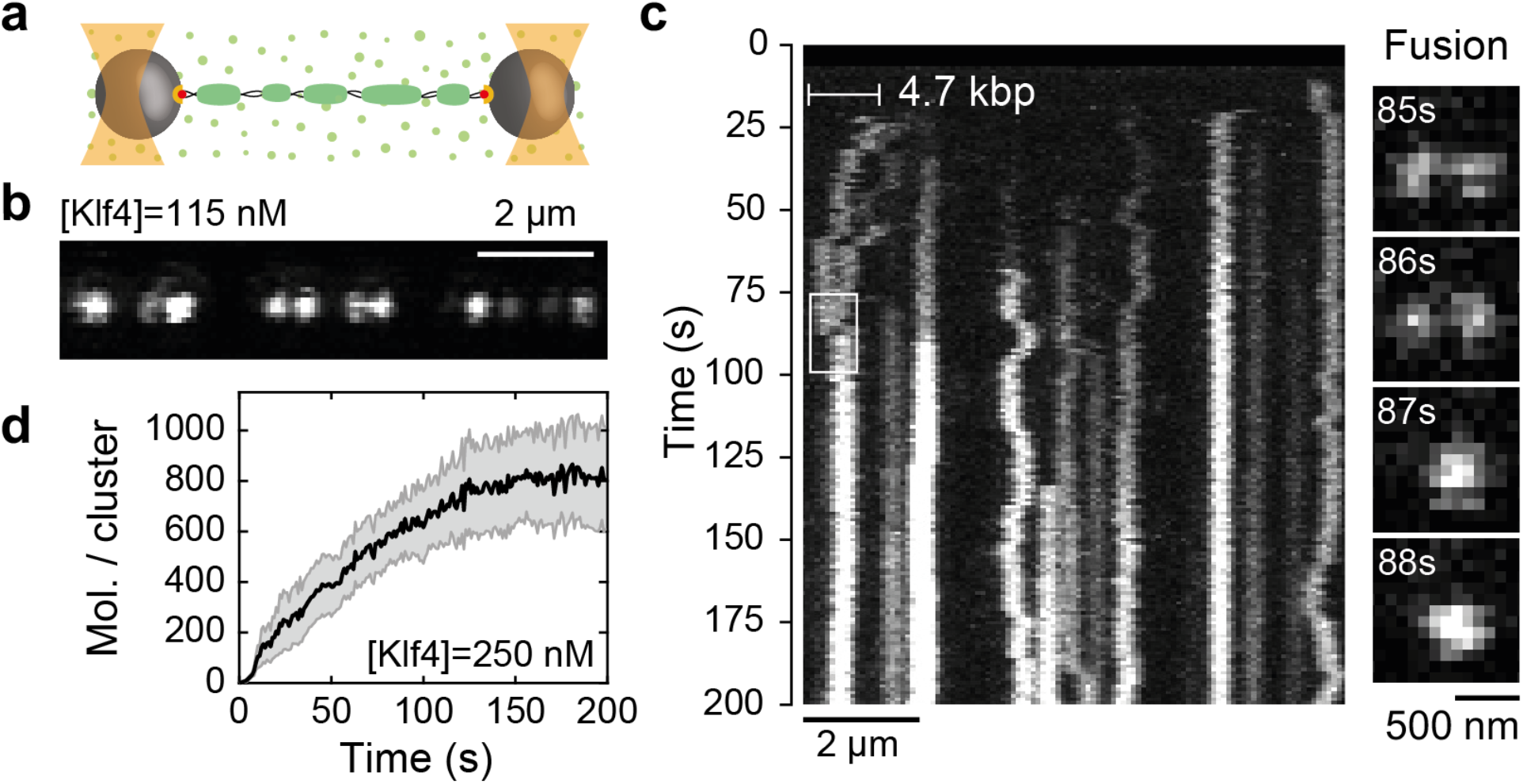
Klf4 forms dynamic foci on a single molecule of λ-DNA stretched in the optical tweezer. **a**, Sketch of the optical tweezer assay depicting a single λ-DNA molecule (black) adhered to a bead at each end via a biotin-streptavidin interaction (red and orange). Beads were held in place in two optical traps (orange cones), and the bead-DNA construct was subjected to approximately 8 pN of tension (Extended Data Fig. 4f, g). Transfer of the construct to a solution containing Klf4-GFP (light green dots) triggered Klf4 foci formation on DNA (dark green). **b**, A representative confocal image of Klf4-GFP on DNA 200 s after exposure of DNA to protein. **c**, Left, kymograph revealing foci formation and dynamics for the experiment shown in **b**. White horizontal bar indicates the approximate extent of foci displacement on DNA (see also Extended Data Fig. 6). Right, foci fusion observed in the indicated region in the kymograph (white box). See also Supplementary Video 1. **d**, The average number of Klf4-GFP molecules per focus increased with time and saturated at approximately 800 molecules. Black line indicates the mean, grey region the standard error of the mean at 95% confidence from 20 foci in 13 experiments at 250 nM Klf4-GFP. Subsequent FRAP experiments displayed nearly complete recovery, indicating exchange of Klf4-GFP molecules between the foci on DNA and solution (Extended Data Fig. 5 and Supplementary Video 2).

To shed light on the physical nature of the Klf4 foci on DNA we investigated their dependence on protein concentration. Fig. 3a shows representative fluorescent images and corresponding traces of Klf4 intensities at different concentrations, recorded 200 s after the Klf4 solution was introduced to the observation chamber. Klf4 foci number and intensity increased with increasing concentration (Fig. 3a). A probability density histogram of the logarithm of pixel intensities at concentrations above 210 nM reveals a bimodal distribution, indicative of two distinct populations of Klf4 foci (Fig. 3b and Extended Data Fig. 9). The peak of the histogram at low intensity characterizes Klf4 regions with, on average, less than one molecule per binding site (corresponding to a 10 bp footprint for binding of the Klf4 Zinc fingers^41,42^; Extended Data Fig. 10a and b). We refer to this mode of association as the adsorbed state. The peak of the histogram at high intensity encompasses Klf4 regions that contain foci with several hundreds and up to a few thousand Klf4 molecules (Extended Data Fig. 10a, c and d). We refer to this mode of association as the condensed state. Intensity histograms as a function of time allow us to visualize the transition from the absorbed to the condensed state, revealing a switch-like behaviour (Fig. 3c): At low Klf4 concentrations (below 80 nM), the histogram is unimodal throughout and rapidly reaches the adsorbed state. At high concentrations (above 210 nM), the histogram first reaches to the adsorbed state, followed by switching to the condensed state at later times. The occurrence of these two states is concentration dependent and shifts from predominantly adsorbed to predominantly condensed with increasing Klf4 concentration (Extended Data Fig. 11). If we define the condensed fraction as the fraction of DNA that is occupied by the condensed state 200 s after exposure of Klf4 to DNA, we observe a sharp increase in the condensed fraction at a concentration C_PW_ of 108 ± 2 nM (Fig. 3d).

**Fig. 3.**
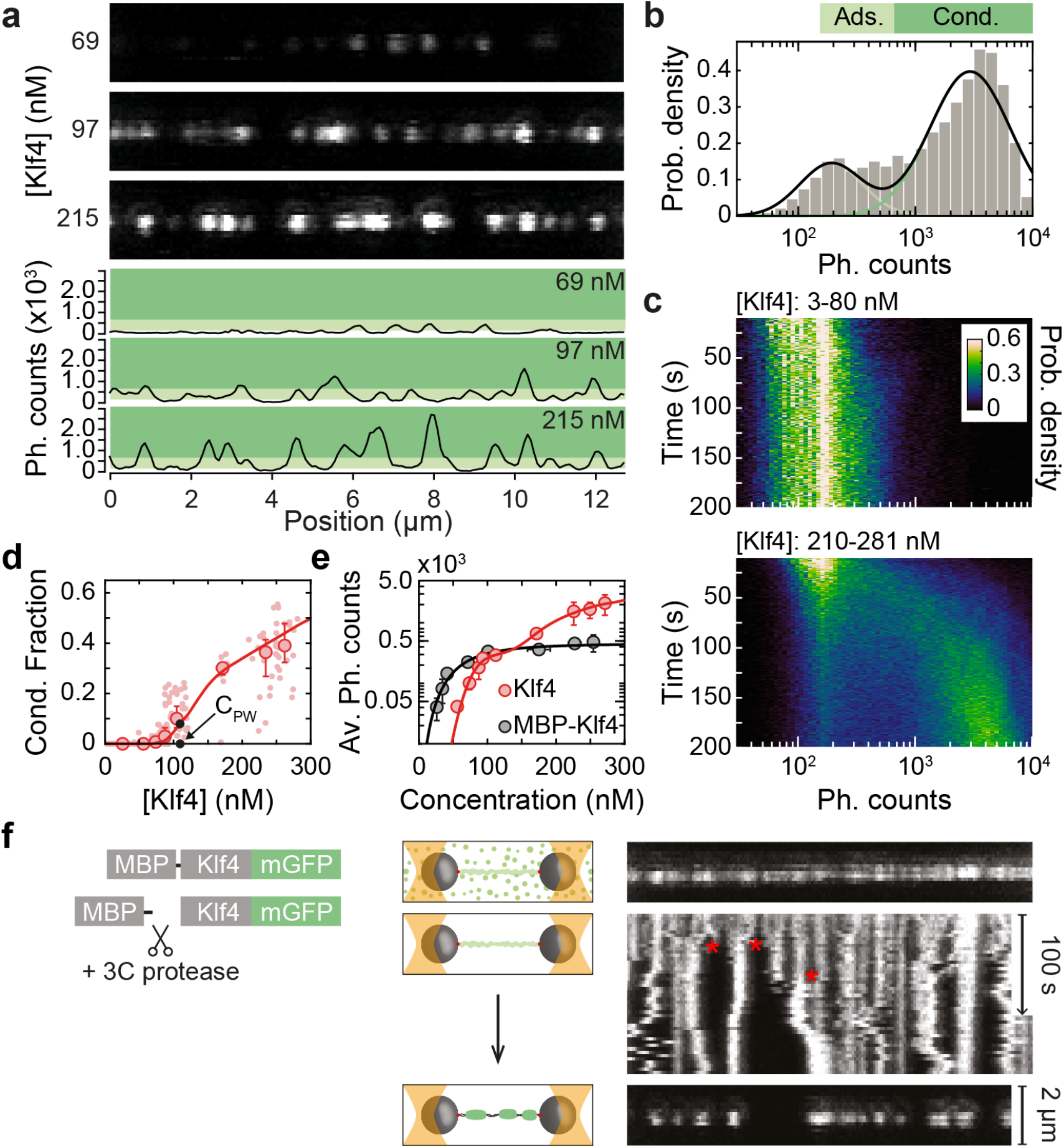
Klf4 foci on DNA switch from an adsorbed to a condensed state. **a**, Representative confocal images at three different concentrations of Klf4-GFP (upper 3 panels) together with corresponding Klf4-GFP intensity profiles (lower 3 panels). The light and dark green areas indicate intensity values that correspond to the adsorbed and condensed state, respectively (as classified in **b**). **b**, A probability density histogram of the logarithm of pixel intensities for all experiments with a Klf4 concentration between 210 and 281 nM (N=37) reveals a bimodal distribution. Black line, fit of the logarithm of intensities to the sum of two normal distributions shown individually by light and dark green lines. The intensity at which these two distributions intersect defines the threshold pixel intensity of 658.5 counts for classification between the adsorbed (light green bar, Ads.) and condensed (dark green bar, Cond.) state (see Methods and Extended Data Fig 9). **c**, Probability density of the logarithm of intensities as a function of time after exposure of DNA to Klf4-GFP for concentrations between 3 and 80 nM (top, N=60) and for concentrations between 210 and 281 nM (bottom, N=37). Experiments in the high concentration range (bottom) reveal switching from the adsorbed to the condensed state with time (see also Extended Data Fig. 11). **d**, Fraction of DNA occupied by the condensed state as a function of Klf4-GFP concentration. Light red dots, individual experiments; red circles, binned medians with error bars denoting the 95% confidence interval as obtained by bootstrapping. Red line, a fit to the heterogeneous two-state model identifies a prewetting concentration of C_PW_ ≈ 108 ± 2 nM. **e**, Mean intensity of Klf4-GFP (red circles) and MBP-Klf4-GFP (black circles) on DNA as a function of concentration. Fits to the heterogeneous two-state model for Klf4-GFP (red line) and a Hill-Langmuir model for MBP-Klf4-GFP are shown (see Supplementary Note). **f**, *In situ* condensation assay. Left, schematic depicting the experimental setup: DNA loaded with MBP-Klf4-GFP containing a 3C cleavage site is transferred to a buffer without MBP-Klf4-GFP but containing the 3C cleavage enzyme. Right, confocal image immediately after transfer (top); Kymograph of the 3C-dependent rearrangement of Klf4-GFP on DNA (middle; red asterisks indicate condensation events); final confocal image after 350 s (bottom). See also Supplementary Video 3 and Extended Data 13.

To trigger the transition from the adsorbed to the condensed state, we took advantage of an observation we made during purification of Klf4: when an MBP-tag (maltose binding protein) was attached to the disordered N-terminus of Klf4 no bulk phase separation was observed (data not shown). When MBP-tagged Klf4 is used in the tweezer assay, a thin adsorption layer forms with, on average, less than one molecule per 10 bp binding site (Extended Data Fig. 12). MBP-tagged Klf4 adsorption to DNA is gradual and follows a Hill-Langmuir isotherm for forming an adsorption layer^43^ (Fig 3e; see Supplementary Note section 4 for a detailed discussion of different binding isotherms). Upon exchange of the buffer to one containing 3C protease without free Klf4 in solution, MBP is cleaved off of Klf4. Strikingly, the adsorbed layer now rearranges into several condensed foci (Fig. 3f, Extended Data Fig. 13 and Supplementary Video 3). We conclude that DNA binding alone is not sufficient for Klf4 condensate formation on DNA, and that the properties of Klf4 that drive bulk phase separation also enable the formation of Klf4 condensates on DNA.

Our results suggest a physical picture where the DNA surface facilitates condensation by locally increasing the density of molecules. Importantly, this condensation happens below the saturation concentration for bulk phase separation. Surface condensation is known to exist in the context of wetting phenomena^44–48^. Here condensates of finite thickness can form on the surface, even at concentrations where the bulk remains well mixed and droplets do not form. Surface condensation is an attractive concept for transcription factors because it provides a mechanism for the formation of small condensates exclusively on DNA that are limited in size by interactions with the DNA surface. This size limitation stems from a balance between surface mediated condensation and bulk mediated dissolution.

Interestingly, our data suggest a switch-like transition from a thin adsorbed to a thick condensed state. In the context of wetting phenomena, such a switch-like behaviour is called a prewetting transition^44–46^ (Fig. 3b-d). Such prewetting transitions occur at a concentration denoted C_PW_, which is below the saturation concentration C_SAT_ in the bulk. Estimates for C_PW_ for Klf4 are shown in Fig. 3d and Table 1. These transitions and their relationship to bulk phase separation are illustrated in Fig. 4a and discussed further in the Supplementary Note. A possible term to distinguish condensates in the bulk from surface condensates is a microphase. Microphases correspond to phase separation that is limited to small scales by the local environment or intrinsic constraints^49^. In our case, the microphase is limited by the requirement of condensation on the DNA surface and is affected by the cylindrical geometry of DNA^10^.

**Fig. 4.**
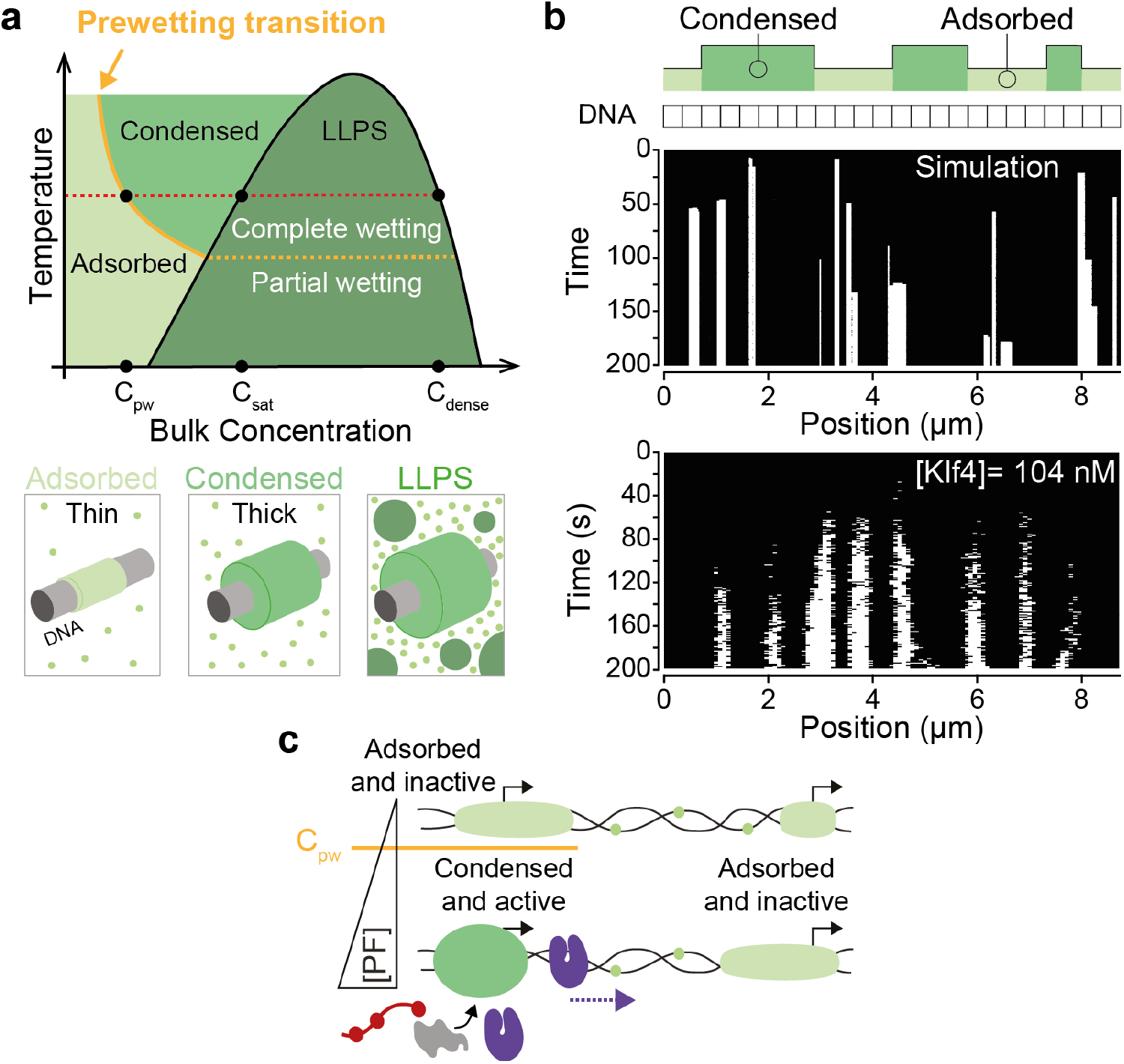
The physics of prewetting captures the switching to a condensed state of Klf4 on DNA. **a**, Phase diagram for a binary fluid in the presence of a surface^10,44^ (top). In the coexistence regime (dark green, liquid-liquid phase separation (LLPS)) and close to the surface, the dense phase transits from partial to complete wetting when the system crosses a characteristic temperature (yellow dashed line). This first-order transition extends into the single-phase region to the left through the prewetting line (solid yellow line). Crossing the prewetting line at a constant temperature (dashed red line) leads to the formation of a condensed thick layer from an initially thin adsorbed layer (bottom row, left and middle). Above the saturation concentration C_SAT_ for LLPS, liquid droplets appear spontaneously in the bulk (bottom row, right). **b**, A heterogeneous one-dimensional two-state model captures the prewetting transition (top, see Supplementary Note). DNA is considered as a 1D lattice of sites which can be in either one of two states, adsorbed or condensed. Sites have an inhomogeneous propensity for condensation drawn from an asymmetric distribution (Extended Data Fig. 14b). An example kymograph reveals that the model captures the spacing, growth, and size of condensed regions of Klf4 (middle) when compared to an experimental kymograph at 104 nM Klf4-GFP and thresholded to display only the condensed state in white (bottom). **c**, We suggest that promoters or enhancers with a more favourable DNA surface for interaction display a lower prewetting concentration C_PW_ for pioneer factors than regions with less favourable surfaces (see text for details). If the pioneer factor concentration is above this threshold, an initially adsorbed state will switch to a condensed state and form a microphase which in turn causes partitioning of downstream factors and the formation of a functional transcriptional hub for activating gene expression.

**Table 1.**
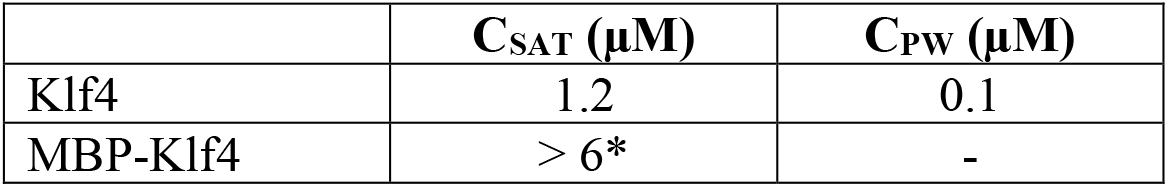
Saturation and prewetting concentrations for Klf4. *Highest concentration tested. Sample does not phase separate.

To examine whether our data are indeed consistent with surface condensation, we put forward a physical theory that considers transitions between thin adsorbed and thick condensed states (Fig. 4b and Supplementary Note). This two-state model is formally equivalent to a random-field Ising model ^50,51^ and describes the statistics for its collective behaviour. We represent the stretched DNA polymer by a one-dimensional set of *N* discrete sites of Klf4 condensation, each of which can be in either a thin adsorbed (*s*_i_=-1) or a thick condensed state (*s*_i_=+1), where *i*=1,..,*N* (Fig. 4b top). At low bulk concentrations the thin adsorbed state is thermodynamically favoured, while at sufficiently high concentrations adsorbed molecules collectively switch to form a thick condensed state. This balance is captured by the energy *h* which is proportional to the bulk chemical potential of Klf4. The effects of DNA sequence on protein binding imply that the DNA is a heterogenous substrate for surface condensation^52^. We capture sequence heterogeneity by introducing a site-dependent bias (*h*_i_) to the chemical potential, which shifts the balance from a thin layer to a condensate (Extended Data Fig. 14b). The free energy is given by

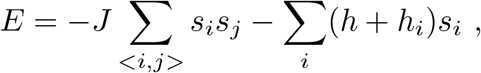

where the first sum is over pairs 〈*ij*〉 of adjacent sites *i* and *j* on the DNA (every pair is counted once). *J* is an energy penalty that is related to the interfacial tension of the surface layer. This coarse-grained and heterogeneous two-state model provides an approach for investigating surface condensation phenomena that is agnostic to geometrical details. Notably, our theory captures key aspects of our experimental data, such as co-existence of thin and thick protein layers, and the switching from the adsorbed to the condensed state upon increasing protein concentration (Fig. 3d, e, 4b and Extended Data Fig. 14).

Taken together, theory and experiment provide strong evidence that Klf4 microphase formation on DNA is an example of a wetting phenomenon below the concentration for bulk phase separation (Fig. 4a). Surface condensates would dissolve if the surface was removed; these condensates are limited in size and are confined to the region where surface interactions allow their formation. A corollary to our observations is that above the bulk saturation concentration, surface condensation would lead to the formation of a thick wetting layer, possibly giving rise to droplet formation via a Plateau-Rayleigh instability^9,53^. This instability governs, for example, dew drop formation on spider webs^54^. While our simplified two-state model describes many aspects of the physics of transcription factor surface condensation, it does not fully consider condensate thickness changes and does not capture the crossover to a Plateau-Rayleigh instability. Future work will be required to expand the two-state model to capture thickness variations of the condensed state.

In this paper we have not investigated sequence dependence of microphase formation on DNA and the role of Klf4 recognition sites. Testing this will require the construction of long pieces of DNA with specific numbers and densities of binding sites, which is technically challenging to achieve. However, we can speculate on the implications of our work for the process by which transcription factors find their target sites. Our observations are consistent with previous suggestions that, at the molecular scale, transcription factors could undergo a 1D diffusive search along DNA, for example in the adsorbed state^55–59^ (Fig. 2c and Extended Data Fig. 6). Specific DNA sequences could trigger the formation of condensed states by locally enabling surface condensation^33^. Furthermore, other surface properties such as DNA methylation or histone modifications could prevent or promote surface condensation, or facilitate or restrict condensate movements^42^. The resulting microphases would then specifically trigger transcription initiation via the recruitment of downstream factors (Fig. 4c). However, such ideas must be tested *in vivo*. For example, it is possible that inside cells, changes in protein levels could shift the system between a surface condensation and a nucleation-growth regime, for instance when cells transform to become cancerous. Such shifts could be tested via controlled increases of protein levels in cells.

Since polymer surface mediated condensation also leads to the formation of liquid-like compartments, features such as fusion and downstream factor concentration that have been observed previously^16,20,22^ are still applicable. The limited size of transcription factor microphases provides a possible explanation of the consistently small size of transcriptional foci *in vivo*^22,60,61^. We suggest that polymer surface mediated condensation provides a general framework to explain the formation of other nuclear condensates such as heterochromatin, paraspeckles or nuclear speckles using methylated chromatin or RNA as a surface, respectively^26,62–66^.

## Methods

### Protein expression and purification

Proteins were expressed in Sf9 cells for 72 h using the baculovirus system^32^. For all Klf4 constructs, the preparation was done on ice using pre-cooled solutions. Cells were resuspended in lysis buffer (50 mM Bis-Tris-Propane pH 9.0, 500 mM KCl, 500 mM Arginine-HCl, 6.25 μM ZnCl2, 5% glycerol, EDTA-free protease inhibitor cocktail set III (Calbiochem), 0.25 U Benzonase (in-house) per ml) and lysed by sonication. The lysate was cleared by centrifugation for 1 h at 13,000 x g and 4°C. The supernatant was incubated with amylose resin (NEB) for at least 30 min at 4°C. After washing with wash buffer I (50 mM Bis-Tris-Propane pH 9.0, 500 mM KCl, 500 mM Arginine-HCl, 5% glycerol), the beads were transferred into Econo-Pac gravity columns (Bio-Rad) and washed with wash buffer II (50 mM Bis-Tris-Propane pH 9.0, 1 M KCl, 500 mM Arginine-HCl, 5% glycerol) followed by wash buffer I. MBP-Klf4 was eluted using elution buffer (50 mM Bis-Tris-Propane pH 9.0, 500 mM KCl, 500 mM Arginine-HCl, 10 mM maltose, 5% glycerol). The eluate was concentrated using VivaSpin 50,000 MWCO concentrators (GE Healthcare or Sartorius) and subjected to size exclusion chromatography at 4°C using a Superdex-200 column (GE Healthcare) and SEC buffer (50 mM Bis-Tris-Propane pH 9.0, 500 mM KCl, 5% glycerol, 1 mM DTT).

After concentrating the sample as described above, the proteins were stored at 4°C for no longer than 2 weeks. MBP-Klf4 was buffer exchanged using Zeba Spin Desalting Columns (Thermo Scientific) into Klf4 buffer (25 mM Tris pH 7.4, 500 mM KCl, 1 mM DTT, 0.1 mg/ml BSA). For MBP-Klf4-GFP and MBP-Klf4-mCherry, unless stated otherwise, the MBP moiety was cleaved off with 10% (v/v) 3C protease (in-house; 1 U/μl) for at least 1 h on ice (see also Extended Data Fig. 1b). For MBP-Klf4-MBP, both MBP tags were cleaved off with 10% v/v 3C protease and 10% (v/v) TEV protease (both in-house) for at least 2 h on ice (see also Extended Data Fig. 3c). In both cases, the sample was spun for 10 min at 20,000 x *g* and 4°C and the concentration was remeasured using either the adsorption at 280 nm or the GFP fluorescence.

### Phase separation assays

Klf4-GFP was kept on ice and diluted with pre-cooled solution to prevent premature phase separation at higher temperatures (Extended Data Fig. 3f). The protein was pre-diluted with cold Klf4 buffer to four times the final concentration and then mixed 1:4 with cold dilution buffer (25 mM Tris pH 7.4, 1 mM DTT, 0.1 mg/ml BSA) to obtain the Klf4 assay buffer (25 mM Tris pH 7.4, 125 mM KCl, 1 mM DTT, 0.1 mg/ml BSA) in a total volume of 20 μl. For assays containing DNA, the dilution buffer would also contain the appropriate amounts of DNA. The samples were mixed by pipetting and 18 μl were transferred into 384 well medium-binding microplates (Greiner bio-one). The samples were incubated at RT for 20 min prior to imaging. Images were taken using an Andor Eclipse Ti inverted Spinning Disc Microscopes with an Andor iXON 897 EMCCD camera and an UPlanSApo 40x/0.95 NA air or a 60x/1.2 NA water-immersion objective (Nikon). Data from at least 3 independent experiments were averaged. Data analysis was performed as described in^67^. For untagged Klf4, differential interference contrast (DIC) microscopy was done using a Zeiss LSM 880 inverted single photon point scanning confocal system utilizing a transmitted light detector and a 40x/1.2 NA C-Apochromat water-immersion objective (Zeiss) which is suitable for DIC.

### Temperature-dependent phase separation

For this, Klf4-GFP was cleaved as described above and the assay was performed as detailed in the section ‘Phase separation assay’ but substituting Tris by 25 mM HEPES pH 7.4 at all steps. A 1 μM solution of Klf4-GFP was mixed with an equal volume of Pico-SurfTM 1 (2% in NovecTM 7500) so that condensates were protected from aberrant surface effects. The sample was imaged on a glass slide which was prepared with thin strips of parafilm to form chambers. A temperature stage was set up as described in^68^. The sample was mounted at 5°C and imaged while the temperature was increased in a step-wise manner as shown in Extended Data Fig. 3f. The imaging was done using a confocal spinning-disc microscope (IX83 microscope, Olympus; W1 SoRa, Yokogawa; Orca Flash 4.0, Hamamatsu) with a 40x air objective (0.95NA, Olympus) controlled via CellSens software.

### Determining the concentration of the dilute phase

The Klf4 samples were set up as for phase separation assays, but instead of transfer to a microplate, samples were incubated for 20 min at room temperature in 1.5 ml Eppendorf tubes. To obtain a standard curve, samples with a final KCl concentration of 500 mM were also prepared and treated in parallel. After incubation, the samples were spun in a temperature-controlled centrifuge at 20,000 x *g* for 15 min at 21°C. 5 μl of the supernatant were added to 15 μl of Klf4 buffer or, in the case of the control samples a corresponding buffer to reach the same final KCl concentration. These samples were transferred to 384 well non-binding microplates (Greiner bio-one) and imaged with a wide-field fluorescence microscope (DeltaVision Elite, AppliedPrecision) using a 10x/0.4 NA dry objective and a Photometrics EMCCD camera. The control samples were used to generate a standard curve that correlates fluorescence intensity with protein concentration. This curve was then used to calculate the original protein concentration in the sample supernatant. Median fluorescence values were obtained using the Fiji software (http://fiji.sc/) and data plotting and fitting was done with R.

### Fluorescence recovery after photobleaching (FRAP)

For Klf4, droplets were essentially prepared as for phase separation assays. However, instead of microplates, glass slides were used. For that, a 1×1 cm square was cut out of a strip of double-sided tape (19 mm x 32.9 mm, Scotch) which was then stuck onto a glass slide. 2 μl of the sample was pipetted onto the glass slide and a PEGylated cover slip was used to seal the chamber in the distance of the tape. The sample was incubated with the cover slip at the bottom for 5 to 10 min at RT before FRAP of settled drops was performed. Photo-bleaching was performed for individual droplets using a 488 nm laser for 2 x 20 ns in an area of approximately 1.5 μm^2^. Recovery of fluorescence intensity within the region of interest was recorded at a rate of 100 ms/frame followed by 500 ms/frame for 50 s each. The pixel resolution was 137 x 137 nm and a 60x/0.9 NA dry objective mounted on a Zeiss Axio Observer.Z1 inverted confocal microscope with a spinning disc and Zeiss AxioCam MRm Monochrome CCD camera were used. Data analysis was performed using the Jython script developed during the Image Processing School Pilsen 2009 which was added to FIJI and is detailed here: https://web.archive.org/web/20200708074426/https://web.archive.org/web/20200504171450/https://imagej.net/

### Electrophoretic mobility shift assays (EMSAs)

Reactions were set-up at 4°C at the indicated final protein concentrations. Klf4 samples contained 25 mM Tris pH 7.4, 125 mM KCl, 6% glycerol, 1 mM DTT, 0.1 mg/ml BSA, 7.5 nM Cy5-dsDNA, 37.5 ng poly-d(IC). Oligonucleotides used can be found in Extended Data Table 1. The absence of condensed material under these conditions was confirmed by fluorescence microscopy (data not shown). The samples were incubated for 20 min at 4°C before they were loaded onto a pre-run 4-20% Novex TBE gel (Invitrogen). Electrophoresis was performed at 200 V for 45 min in TBE buffer (89 mM Tris, 89 mM boric acid, 2 mM EDTA). The gels were then imaged using a Typhoon 9500 FLA fluorescent imager (GE Healthcare). Band intensity was determined using the Fiji software (http://fiji.sc/) and data plotting and fitting was done with Matlab. The following expression was used to fit the data^69^:

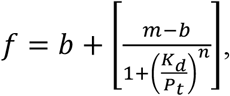

where *P_t_* is the total protein concentration and *K_d_* is the dissociation constant. m and b are normalization factors for the upper and lower asymptotes of the DNA titration curve and n is the Hill coefficient.

### Hardening assay

Because Klf4-mCherry does not phase separate at pH 7.4 (data not shown), a lower pH buffer was used than for all other assays. The proteins were buffer exchanged into 25 mM Tris pH 7.4, 250 mM KCl, 1 mM DTT, 0.1 mg/ml BSA and cleaved as described above. For the phase separation assay, the protein was diluted with the above buffer to 8 μM and mixed with an equal volume of 25 mM PIPES pH 6.5, 1 mM DTT, 0.1 mg/ml BSA to obtain a final Klf4 concentration of 4 μM. The sample was transferred into a 384-well microplate, incubated for 20 min and imaged as described in the section ‘Phase separation assay’. Phase separation of both, Klf4-GFP and Klf4-mCherry as well as an equimolar mixture of both was confirmed as shown in Extended Data Fig. 8. For the hardening assay, 4 μM of Klf4-GFP were phase separated and incubated for 50 min before imaging. Next, an equal volume of freshly diluted Klf4-mCherry was added to obtain a final concentration of 2 μM for both proteins in the same conditions as described above (50% pH 7.4/50% pH 6.5). The mixture was imaged after 1 min and 10 min in the same way as before.

### dCas9-EGFP λ-DNA labelling

Recombinantly expressed dCas9-EGFP was stored at 5.3 mg/ml at −80°C in storage buffer (250 mM HEPES pH 7.3, 250 mM KCl) and thawed 1 h prior to the experiment. sgRNAs were designed against four adjacent target loci on λ-DNA (Extended Data Table 2). The spacing between adjacent target sequences was adjusted to 40-50 bp to prevent steric hindering of adjacent dCas9-sgRNA complexes. Guide RNAs were expressed and purified using commercial kits (MEGAscript T7 Transcription Kit, Invitrogen and mirVana miRNA isolation Kit, Invitrogen); stored in ddH2O at 0.6-1 mg/ml at −80°C and thawed 1 h prior to the experiment. We first mixed 2 μl of dCas9-EGFP with 38 μl of complex buffer (20 mM Tris-HCl pH7.5, 200 mM KCl, 5 mM MgCl2, 1 mM DTT) and then 5 μl of this 20x dCas9-EGFP dilution with 4 μl sgRNA stock which contained all four sgRNAs in equal stoichiometries. The reaction volume was adjusted with complex buffer to 50 μl and incubated at room temperature (22°C) for 30 min. After incubation, we mixed 10 μl of the dCas9-sgRNA complex reaction with 1 μl of 5 nM biotinylated λ-DNA. The reaction volume was adjusted with reaction buffer (40 mM Tris-HCl pH 7.5, 200 mM KCl, 1 mg/ml BSA, 1 mM MgCl2 and 1 mM DTT) to 50 μl. The sample was incubated at room temperature (22°C) for 30 min. The final dCas9-sgRNA-λ-DNA complex was then diluted to 5 pM in the corresponding assay buffer.

### Optical tweezers with confocal microscopy

Optical tweezer experiments were performed on a Lumicks C-trap instrument (Amsterdam, Netherlands) with integrated confocal microscopy and microfluidics. Bacteriophage λ-DNA was biotinylated on both ends as described elsewhere^38^. Attachment of λ-DNA-dCas9 complex to 4.42 μm Spherotec streptavidin coated polystyrene beads was done using the laminar flow. For all experiments, the trap position was kept constant to render an average force of 8.22 +/- 2.65 pN (Extended Data Fig. 4).

The protein stock was centrifuged for 10 min at 20,000 x *g*. The supernatant concentration was measured and diluted in the phase separation assay buffer following a dilution series. A solution containing the maximum concentration of a given series was flushed into the flow cell. After recording of 10 to 15 experiments, the remaining volume in the syringe was removed, the protein diluted and reloaded into the syringe. The flow chamber was flushed prior to each experiment and sealed during the course of it.

For confocal imaging, the 488 nm laser was used for excitation, with emission detected in the channel with blue filter 525/25 nm. After a λ-DNA molecule had been tethered between the beads, an image of the dCas9-EGFP probe was acquired in the buffer channel with a 10% excitation intensity (we refer to this imaging setting as high excitation). We then started continuous acquisition with 5% excitation intensity while the beads-DNA system was transferred to the channel of the microfluidics chip containing the protein of interest. The interaction process was monitored for 200 s at a frame rate of ~1/s with a low pixel integration time of 0.08 ms (Fig. 2c and Extended Data Fig. 6). After 200 s, an image was acquire using the high excitation imaging conditions. Analysis of the intensity distributions and quantification of number of molecules (Fig. 3a, b and Extended Data Fig. 9, 10 and 12) was done using high excitation settings. In order to determine the number of molecules per cluster over time (Fig 2d), time series were acquired using a pixel integration time of 1 ms (referred to as low excitation, see Extended Data Fig. 4), conditions in which the dCas9-EGFP probe was detectable for the first few frames, before the beads-DNA system reached the protein solution.

For the *in situ* condensation assay (Fig. 3f and Extended Data Fig. 13), after the binding process had been recorded, the beads-DNA system was transferred back to the buffer channel containing either the assay buffer or the assay buffer with 2% (v/v) 3C protease (in house, 1 U/μl). This process was recorded for more than 500 s with low excitation settings at a frame rate of 0.2/s.

FRAP experiments were performed as follow: after a binding experiment (low excitation settings), the chamber was gently flushed. A pre-bleach time series was acquired for 20 s. A smaller ROI (FRAP ROI) was imaged with high excitation laser intensity (90% excitation). To capture recovery, a 200 s long time series was then acquired at a frame rate of ~1/s (Extended Data Fig. 5).

## Data analysis

### Background subtraction and concentration calibration

We selected an ROI of 15×60 and 4×40 (vertical x horizontal directions) pixels for high and low intensity images respectively, centred in the middle of the confocal image in the horizontal direction and spanning from the top. For background subtraction, the average intensity in this window was obtained and subtracted from each pixel in an image. For time series, the average background intensity of the last 10 frames was subtracted.

To determine the concentration of individual experiments, a calibration curve of background intensity vs concentration was built for each dilution series using the last 3 experiments of each dilution step. The background intensity for each experiment was then converted to concentration units using the formula:

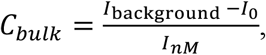

where *I*_background_ is the mean intensity of the background; *I_nM_* is the slope of the calibration curve and *I_o_* the intercept (Extended Data Fig. 7 and Extended Data Table 3).

### Intensity emission per EGFP

For each experiment, confocal images of the dCas9-sgRNA-λ-DNA complex were acquired using high excitation settings. For time series (low excitation settings) the dCas9 probe was detectable for the first few frames, before the beads-DNA complex reached the protein solution. To confirm the position of the dCas9 probe, intensity profiles along the DNA were aligned using the beads centre as a reference and flipped when required. To confirm the position of the target sequence, the target locations were superimposed with the average profile (Extended Data Fig. 4). Sequence information was converted to spatial units by taking into account the extension per base pair (xbp = 0.32 nm/bp) at the average experimental force. The integration of the total intensity in a ROI of 21 x 21 pixels around the detected probe rendered the total number of counts under a given imaging conditions. The probability distribution of integrated counts for several experiments exhibits a multi-mode gaussian distribution, consistent with having 4 sites for dCas9 binding in λ-DNA (see Extended Data Table 2). A fit to a gaussian mixture model rendered the mean and standard deviation of each mode. The emission intensity per EGFP was then calculated as:

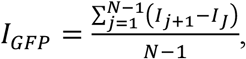

where *I_j_* is the mean of mode j and *N* is the number of modes (See Extended Data Fig. 4 and Extended Data Table 4).

### Cluster trajectories and number of molecules over time

Identification of cluster trajectories was done using the Python package Trackpy^70^. In brief, in each frame the algorithm finds particles based on a comparison between the local pixel intensity and the pixel intensity in the surrounding area with radius PR. Particles in sequential frames that are within the search radius (SR) and persist for a number of frames t_memory_ as imposed by the user are linked together to generate a trace. In a final step, traces might be accepted or discarded based on the total duration t_min_ (Extended Data Fig. 6). The number of molecules per cluster over time (Fig. 2c) was determined by dividing the integrated intensity of a particle (‘mass’ in the Trackpy output) by the intensity per EGFP. Position of clusters along the DNA in base pair units (Fig. 2c and Extended Data Fig. 6) was calculated as: Position = (X_px_ * s_p_) / x_bp_, where X_px_ is the position in pixel units and s_p_ = 80 nm is the pixel size.

### Intensity distributions

Pixel values used in the calculation of the intensity distributions were obtained as follows: after background subtraction (to remove the contributions from the protein in solution), the maximum projection intensity profile along the DNA was determined in a region of 20 pixels around the DNA axis (see Extended Data Fig. 9). We next filtered the profiles with a spatial mask. In brief, using the findpeaks algorithm from Matlab we detected the peaks above a threshold corresponding to the background value of the background subtracted image (see Extended Data Fig. 9). Data points in a 5 pixels window, along the horizontal direction and centred at the position of each peak were selected. The window was displaced from left to right and accepted if there was no overlap. From the histograms of pixel intensity values obtained, we computed the probability density of the logarithm of pixel intensities^71^. The probability density vs the logarithm of intensities was fitted to either one or two components gaussian mixture model in linear scale. To compare the intensity distributions (Fig. 3b) with the intensity distributions over time (Fig. 3c and Extended Data Fig. 11), time series images were multiplied by a factor (13.4 ± 2.9) to compensate for the intensity value differences between low and high excitation imaging conditions. From the last frame of each time series and the corresponding high excitation image acquired immediately after, we computed the mean intensity in a ROI of 30×100 pixels in the centre of the confocal image. Time series intensities were multiplied by the ratio of these means.

### Kymographs

Kymographs shown throughout this work were built using the maximum projection intensity profiles.

### Classification of pixels into adsorbed or condensed

Pixels were classified into adsorbed or condensed based on their intensity. An intensity above the background and below the layer threshold resulted in a classification as adsorbed while an intensity above the layer threshold was classified as condensed. To determine the background threshold, we extracted the background values (after background subtraction) along a line away from the DNA and pulled together all the experiments corresponding to Klf4-GFP. The probability density of the logarithm of pixel intensities was fitted to a normal probability density function. The background threshold was defined as the mean plus 3 times the standard deviation of this distribution (Extended Data Fig. 9). To determine the layer threshold, we computed the probability density of the logarithm of pixel intensities along the masked maximum projection profiles pulling together 60 Klf4-GFP experiments recorded at low concentrations ([Klf4]: 3-80 nM). We extracted the mean value of this distribution by fitting to a normal probability density function. We next computed the same quantity for 37 experiments recorded at high concentrations ([Klf4]: 210-281 nM). Here, the probability density shows bimodality and we fitted this distribution to a two components gaussian mixture model, constraining the mean of the low intensity mode to the value obtained at low concentrations. Fitting was done in Matlab using the *NonlinearLeastSquares* method and weights of 1/(Probailty density)+w, where w=10 sets the strength of the weights.

### Condensed fraction

The condensed fraction (Fig. 3d and Extended Data Fig. 14) was determined for each experiment as the number of pixels falling into the condensed category divided by the length of the considered ROI (161 pixels). For this calculation, we considered the pixels obtained by the masking procedure (see section Intensity distributions and Extended Data Fig. 9). Binned medians with error bars (95% confidence interval) were obtained by bootstrapping (*bootci* function from Matlab) using 10000 bootstrap samples. Binned medians contain 11 to 36 individual experiments.

### Number of molecules per binding site

The number of molecules per binding site was computed as Molecules/binding 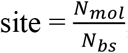, where, N_bs_ = L_bp_/l_bs_ is the number of binding sites of length lbs (assumed to be l_bs_ = 10 bp throughout this work^41,42^), L_bp_ = L_px_ * s_p_ / x_bp_ is the length of the considered region in base pair units and Nmol is the number of molecules in the considered region. To determine the number of molecules in the adsorbed (condensed) state, we first determined the pixels falling into the adsorbed (condensed) classification. Then, we integrated the intensities along the direction perpendicular to the DNA generating an integrated profile with the same spatial dimension as the maximum projection profile (used for pixel classification). Next, we integrate over all the pixels contained in the adsorbed (condensed) state and divide this quantity by the intensity per GFP to compute the number of molecules. The number of binding sites contained in the adsorbed (condensed) regions was determined as the number of pixels classified as adsorbed (condensed) multiplied by the appropriate constants.

## Acknowledgments

S.W. and A.K. were supported by EMBO Long-Term Fellowships (ALTF 708-2017 and ALTF 1069-2017, respectively) and S.G. by the ELBE fellowship program and the Max-Planck-Gesellschaft. This project has received funding from the European Union’s Horizon 2020 research and innovation programme under the Marie Sklodowska-Curie grant agreements No 791147 and No 798297. A.A.H acknowledges support from the MaxSynBio Consortium and the NOMIS foundation. S.W.G. was supported by the DFG (SPP 1782, GSC 97, GR 3271/2, GR 3271/3, GR 3271/4) and the European Research Council (grant 742712). We thank the staff and students of the 2018 and 2019 MBL Physiology courses where this work was started, in particular P. Sil, A. Fenix, A. De Simone, A. A. Bhat, S. Lembo, J. Lippincott-Schwartz, R. Phillips, D. Fletcher, N. King, C. Ott and J. Brzostowski. We thank Andrei Pozniakovsky for help with cloning and Anatol W. Fritsch for assistance with undertaking spinning disk confocal microscopy experiments with the Olympus IXplore IX83 microscope. Further, we wish to thank the following MPI-CBG facilities for support with this project: the Light Microscopy Facility, Benoit Lombardot and the Scientific Computing Facility and the Technology Development Studio. We also thank J. Kondev, I. Cisse, P. Tomancak and J. Howard for discussions and comments on the manuscript.

## Supplementary Note

### 1 Two-state model for surface condensation of pioneer factors on DNA

Here we develop a physical theory of surface condensation of pioneer factor protein Klf4 on DNA. Klf4 exhibits bimodal distribution of intensities along the DNA; the lower and higher intensity modes correspond to a thin adsorbed and a thick condensed state, respectively (see Fig. 3a-d). Motivated by these observations we build a two-state model which is formally equivalent to an Ising model on a linear lattice^72^. In this model (Fig. 4d), the stretched DNA is represented by a one-dimensional set of *N* discrete sites. Each of these sites can either be in state *s_i_* = –1 corresponding to the adsorbed state, or in state *s_i_* = +1 that corresponds to the condensed state, where *i* = 1,.., *N*. The coarse-grained free energy *E* for the model is given by

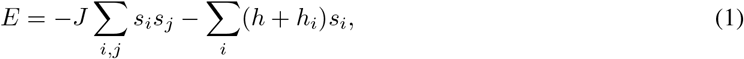

where *J* quantifies the nearest neighbor interactions between the states. Note that in the present context of surface condensation *J* is related to the interfacial tension of the surface layer and *h* is proportional to the bulk chemical potential of the proteins. Proteins bind to different parts of the DNA with different binding affinities^52^. To take into account DNA sequence heterogeneity we introduce a site-specific random bias *h_i_*, where 2*h_i_* is the free energy difference between the −1 and +1 states. This model is formally equivalent to the Random field Ising model (RFIM)^50,51^. For a dilute solution, *h* can be written as^58^

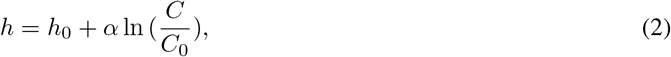

where *C* is the bulk protein concentration and *α* is a constant with the same dimension as *h. h*_0_ is proportional to the chemical potential at reference concentration *C*_0_. In what follows we use *h*_0_ = 0. The partition function for the system is given by

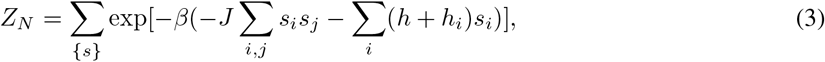

where 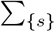 denotes the sum over all possible configurations *s_i_*, and *β* = 1/*k_B_T*. All the relevant equilibrium properties of the system can be computed from the partition function.

#### 1.1 Condensation as a function of bulk protein concentration

In order to compare our model with experimental data we calculate the condensed fraction *ϕ*, defined as the fraction of DNA length occupied by the condensed state as a function of the bulk protein concentration. Two scenarios are considered, namely the i) homogeneous model where the site-specific random bias *h_i_* = 0, and the ii) heterogeneous model for which *h_i_* ≠ 0.

##### 1.1.1 Homogeneous model

We employ the transfer matrix method to obtain the partition function^72^ for the homogeneous model, which in the large *N* limit is given by

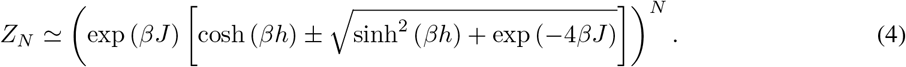

Using the partition function, the total free energy *F* of the system can be computed from 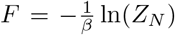. To obtain the condensed fraction *ϕ*, we calculate the average state occupancy per site 〈*s_i_*〉 = *∂F*/(*N∂h*). Hence, the condensed fraction is given by

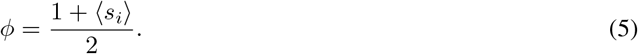

From Eqs.4 and 5, we obtain the following expression for the condensed fraction

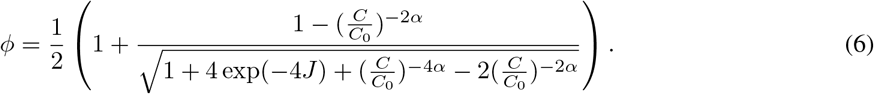

In the ensuing sections we use this expression to compare the homogeneous model to the data shown in Fig. 3d and Extended Data Fig. 14a.

##### 1.1.2 Heterogeneous model

Next, we consider the heterogeneous model with non-vanishing *h_i_*. To compute the condensed fraction (*ϕ*), values for site-specific random bias are drawn from a Gamma distribution which is defined as

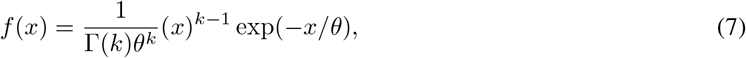

where *x* > 0, *k* and *θ* are shape and scale parameters respectively. The range of Gamma distribution is 0 to ∞. Owing to our sign convention, we take random values x from the gamma distribution with *h_i_* = –*x* for the *i*-th site. A negative *h_i_* implies that the *i*-th site is more likely to be in the −1 state than in the +1 state. For a given realization of *h_i_*, we compute the corresponding transfer matrices along the DNA. To calculate the partition function, we calculate the product of these transfer matrices numerically using MATLAB. From the partition function, we extract the condensed fraction (*ϕ*) by following the procedure outlined in section 1.1.1 above.

#### 1.2 Dynamics of condensation

To compare our model to experimental kymographs (Fig. 2c), we define a kinetic formulation for representation of the model that allows for studying the dynamics of protein condensation on DNA. We start with the coarsegrained free energy of the model defined in Eq.1. This expression is used to define the switching rates between the absorbed and condensed states using the detailed balance condition as described in the next section.

##### 1.2.1 Usage of the detailed balance condition to define state-switching rates

To understand the dynamics of condensation, we study how the system evolves to equilibrium by employing the detailed balance condition, which ensures consistency with equilibrium. The detailed balance condition is given by

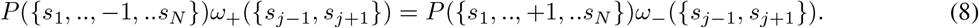

Here *P*({*s*_1_,.., −1,.. *s_N_*}), and *P*({*s*_1;_+1,.. *s_N_*}) are the probabilities of having the *j*-th site to be either in the −1 or +1 state, while states of the other sites are given by *s*_1_, *s*_2_,.. *s_N_* respectively. *ω*_+_({*s*_*j*–1_, *s*_*j*+1_}) is the rate of transition of *s_j_* from the −1 to +1 state, while *ω*_–_({*s*_*j*–1_, *s*_*j*+1_}) is the rate of transition from the +1 to −1 state. To define the rates of state-flipping, we consider the effective field on a given site due to its neighbors. For instance, for the *j*-th site the effective field is given by

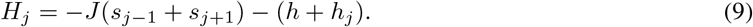

The ratio of switching rates for the *j*-th site (using Eqs.8 and 9) then is

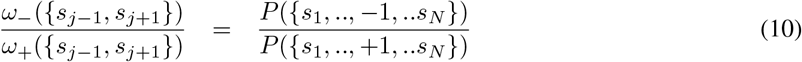

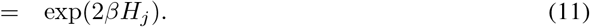

We choose the following form for the rates

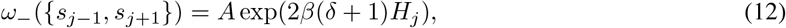

and

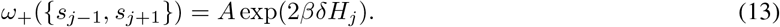

*A* and *δ* are characterized by the timescales of transition^73^ between adsorbed and condensed states in the system. Evidently, these rates satisfy the detailed balance condition. By utilizing these rates, we obtain model kymographs from simulations, which can be compared with data.

### 2 Comparison of model results with experimental data

We investigate if the two-state model can account for our measurements of the condensed fraction as a function of the bulk protein concentration (Fig. 3d), the average intensity along DNA (Fig. 3e), and dynamics of condensation (Fig. 4b).

#### 2.1 Homogeneous two-state model

First, we consider the homogeneous model to fit the condensed fraction data shown in (Fig. 3d). Using Eq.6, we fit condensed fraction for Klf4 as a function of the bulk protein concentration, as shown in Extended Data Fig. 14a. The best-fit parameter values are *J* = 3.37 kT, *C*_0_ = 292 nM, and *α* = 0.0016 kT. Note that the homogeneous model cannot account for the switch-like behavior from adsorbed to condensed layer.

Next, we compare kymographs, as obtained from the homogeneous model and experiments. To generate the kymographs, we choose *N* = 5000 as the number of sites on the DNA. Initially all the sites are chosen to be in the —1 state (adsorbed state). Starting from this state configuration, we observe how the system evolves to condensation as the bulk concentration in increased. We use the parameter values obtained from the condensed fraction fit to simulate the time evolution of the system. A representative kymograph corresponding to a concentration of *C* = 120 nM is shown in Extended Data Fig. 14c. It reveals a pattern of condensed states of variable size. Furthermore, clusters of condensed states emerge and dissolve as a function of time. The details of the simulations are discussed in the section 3.

#### 2.2 Heterogeneous two-state model

To study if the heterogeneous model can capture the data observations, we fit this model to condensed fraction data (Fig. 3d and Extended Data Fig. 14a). We extract the following parameter values: *J* = 6 kT, *C*_0_ = 35 nM, *α* = 0.5 kT, shape parameter *k* = 0.1 (dimensionless), and the scale parameter *θ* =18 kT.

To define the prewetting concentration, we find the point of maximum slope on the curve defining the condensed fraction as a function of bulk protein concentration (see Fig. 3d). The concentration corresponding to this point is defined as the characteristic prewetting concentration *C*_PW_. For Klf4, *C*_PW_ ≈ 108 ± 2 nM.

Next, we generate kymographs for this model using the fitted parameter values. An example kymograph (see Extended Data Fig. 14d) of the heterogeneous model corresponding to a concentration of *C* = 120 nM is shown. It exhibits clusters of condensed states of various sizes, distributed randomly along the DNA. These clusters, once emerged, remain on the DNA for the observed simulation time window. Furthermore, the clusters are pinned to regions of DNA.

##### 2.2.1 Fit to the mean Klf4 intensity along DNA

The mean intensity along the DNA (Fig. 3e) was calculated as the mean value of all the pixels in an intensity profile along the DNA. In the case of MBP-Klf4, the data were fitted to a Hill function:

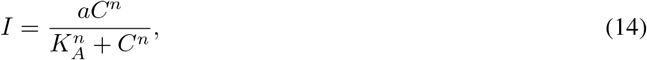

where *C* is the bulk protein concentration. The fit parameters are defined as follows: *K_A_* = 70.0 nM is the protein concentration when the intensity value is half the maximum, *n* = 2.1 is the Hill coefficient, and *a* = 457.8 is the proportionality constant between the fraction of protein bound to the DNA and the corresponding intensity.

The mean intensity of Klf4 along the DNA first increases as a function of the bulk protein concentration and saturates to a level comparable to MBP-Klf4 (~ 400). As we further ramp up the bulk concentration, the mean intensity rises again, showing a deviation from a pure Hill-Langmuir behavior. To capture this non-monotonic trend, we need to account for the dimensionality of the adsorbed and condensed states, which remains absent in the two-state model. Hence we fit the data to the heterogeneous Ising model, convoluted with Hill functions. The fit function is defined as:

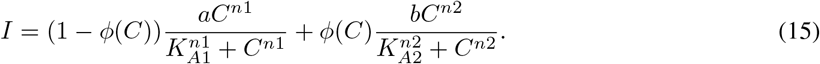

Here *ϕ*(*C*) is extracted from a fit of the two-state model to the condensed fraction data in Fig. 3d, and for each concentration is imposed as a fixed parameter. The remaining six parameters in Eq. 15 are extracted from the fit. Since for *ϕ* = 0, the average intensity results from the adsorbed state exclusively, *K*_*A*1_ = 83.2 nM, n1 = 6.4, and *a* = 355 have a similar interpretation as in Eq. 14 for MBP-Klf4 binding. As the bulk protein concentration is increased further, the condensed state emerges, contributing to the rise in mean Klf4 intensity. To capture this rise we introduce the second part of Eq. 15. For *ϕ* =1, the mean intensity of Klf4 results from the condensed state only, and saturates at a value characterized by *b* = 4461.4. *K*_*A*2_ = 192.8 nM is the protein concentration when the Klf4 intensity value is half the saturation level, n2 = 6.3 is the Hill coefficient for the growth of the condensed state.

### 3 Simulation method for obtaining model kymographs

To generate kymographs for the two-state model, we choose *N* = 5000 as the number of sites on the DNA. Hence, each site in our model accounts for approximately one Klf4 binding site (a single Klf4 molecule occupies about 10 bp of DNA^41,42^). The λ-DNA used in our experiments is about 48 kbs long Eqs. 12, and 13 are used to define the rates of state-switching. We choose *A* = 2000/unit time, and *δ* = 0.5 for the rates. The choice of *A* and *δ* alters the timescale of condensation. The values we choose are consistent with the onset of microphase formation in experiments, which is about 10 – 20 seconds. *β* is set to unity.

We employed the Gillespie algorithm^74^ to generate the kymographs, using the rates defined in Eqs. 12, and 13. A single time-step of simulations is performed as follows: one of the set of all possible reactions is chosen according to its relative weight, and the state of the system is updated appropriately. At the same time, the time elapsed since the last step is chosen from an exponential distribution, whose rate parameter equals the sum of rate parameters of all reactions. We iteratively repeat this process until the system reaches steady-state, whereby the number distribution of +1 states in the system remains unchanged.

### 4 Description of Langmuir and BET isotherms

Single-layer adsorption can be characterized by a Langmuir isotherm^43^. We have system consisting of a surface and protein molecules that bind to the surface. Let’s assume that the total area of the surface is *A*, whereas each molecule occupies an area of *a*^2^. Hence the number of putative protein binding sites on the surface is *N_tot_* = *A*/*a*^2^. A protein binds to a given binding site with binding energy *ϵ*, and the bulk chemical potential is *μ*. If the binding of protein molecules to the surface is not cooperative, binding to every site on is independent. There are two possible configurations for every binding site: it is either occupied or unoccupied. The partition function per site is then given by a product from independent contributions of binding sites

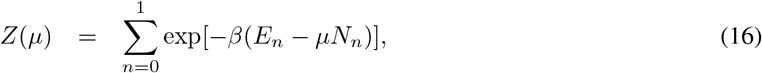

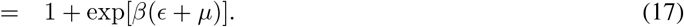

Here the binding site has energy *E_n_* = *ϵ* when the site is occupied by a protein molecule and *E_n_* = 0 when the site is unoccupied. *N_n_* = 0 if the site is unoccupied, *N_n_* = 1 if it’s occupied. Using Eq. 17, we obtain the grand potential

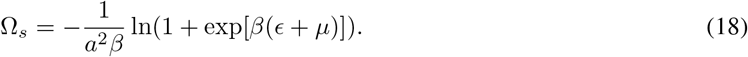

The surface concentration of protein is given *∂*Ω_*s*_/*∂μ* = –*C_s_*. From Eq. 18, we get an expression for *C_s_* as a function of the bulk chemical potential, which is given by

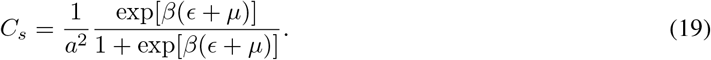

A BET Isotherm generalizes the concept of Langmuir isotherm for multi layer adsorption^75^. In addition to single molecule binding to the surface (with binding energy *ϵ*), a protein molecule can attach to another protein molecule bound to the surface (with binding energy *E*). This way multiple molecule layers can form on the surface. The partition function per site is given by

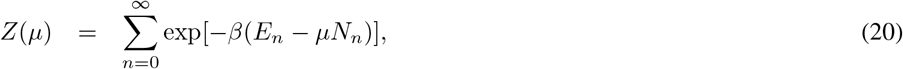

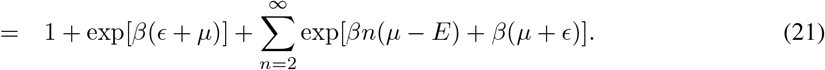

*E_n_* = *ϵ* when a site on the surface is occupied. For the subsequent layers, the binding energy is *E_n_* = *E*. *n* is the layer number.

We now define *x* = exp[*β*(*μ* – *E*)], and *y* = exp[*β*(*μ* + *ϵ*)] = *cx*, such that *c* = exp[*β*(*E* + *ϵ*)]. From Eq.21, we have

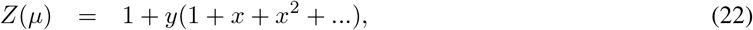

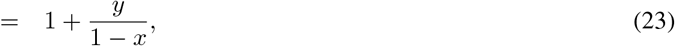

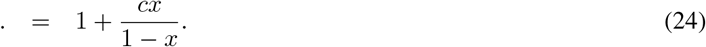

The grand potential is given by

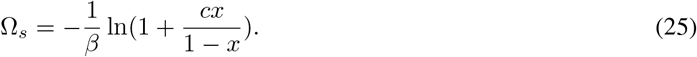

Since *∂*Ω_*s*_/*∂μ* = –*C_s_*, we get

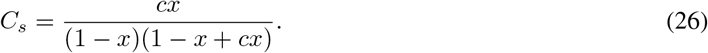

This is the expression for *C_s_* as a function of^76^ *x* = exp[*β*(*μ* – *E*)].

## Supplementary Video Legends 1-3

**Video 1**

Representative example of Klf4 binding to λ-DNA (same data as shown in Fig. 2b and c). The movie starts before the beads-DNA complex reaches the protein solution, at which point there is an increase in the background intensity. For this representation, background intensity due to the protein solution was not subtracted.

**Video 2**

Representative example of a Klf4 FRAP experiment (same data as shown in Extended Data Fig. 5). Red box shows the photo-bleached region. For this representation, background intensity due to the protein solution was not subtracted.

**Video 3**

Representative example of the *in situ* condensation assay (same data as shown in Fig. 3f). The movie starts while the beads-DNA complex is in the MBP-Klf4 protein solution. After a few frames, the complex is transferred to a buffer containing the 3C cleavage enzyme evidenced by a drop in background intensity. For this representation, background intensity due to the protein solution was not subtracted.

## Extended Data Figures 1–14

**Extended Data Fig. 1:**
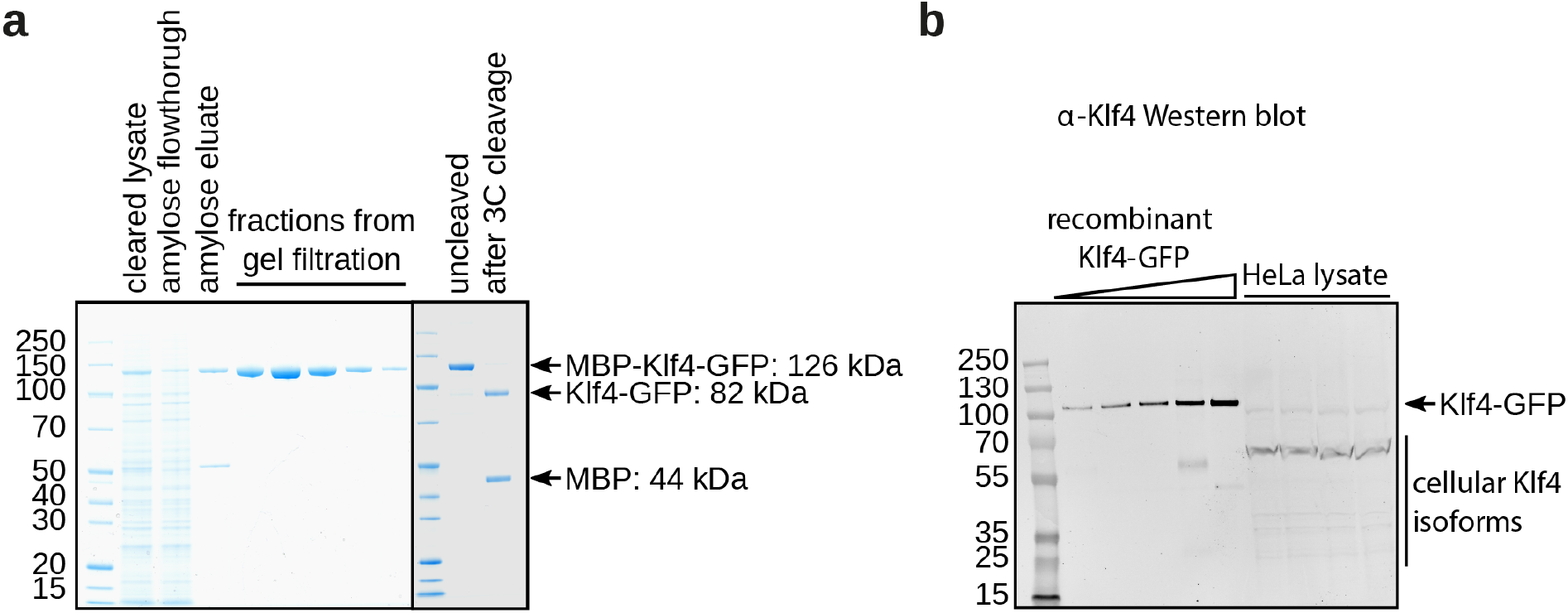
Klf4 can be recombinantly purified and quantified in cell lysates. **a**, SDS gel showing a representative purification of MBP-Klf4-GFP. First, the Sf9 insect cell lysate is cleared by centrifugation (cleared lysate) before it is subjected to amylose resin (amylose flowthrough and amylose eluate). The eluate is then concentrated and further purified by size exclusion chromatography (fractions from gel filtration). Right before the assay, the concentrated MBP-Klf4-GFP sample (uncleaved) is treated with 3C PreScission protease (see Methods) to remove the MBP-tag from Klf4-GFP (after 3C cleavage). **b**, The concentration of intracellular Klf4 was estimated using HeLa lysates and Western blotting with fluorescent secondary antibodies (see Methods). A representative example blot is shown. Recombinantly expressed and purified Klf4 was used to generate a standard curve on each Western blot. With this, the amount of Klf4 in HeLa lysates was determined.

**Extended Data Fig. 2:**
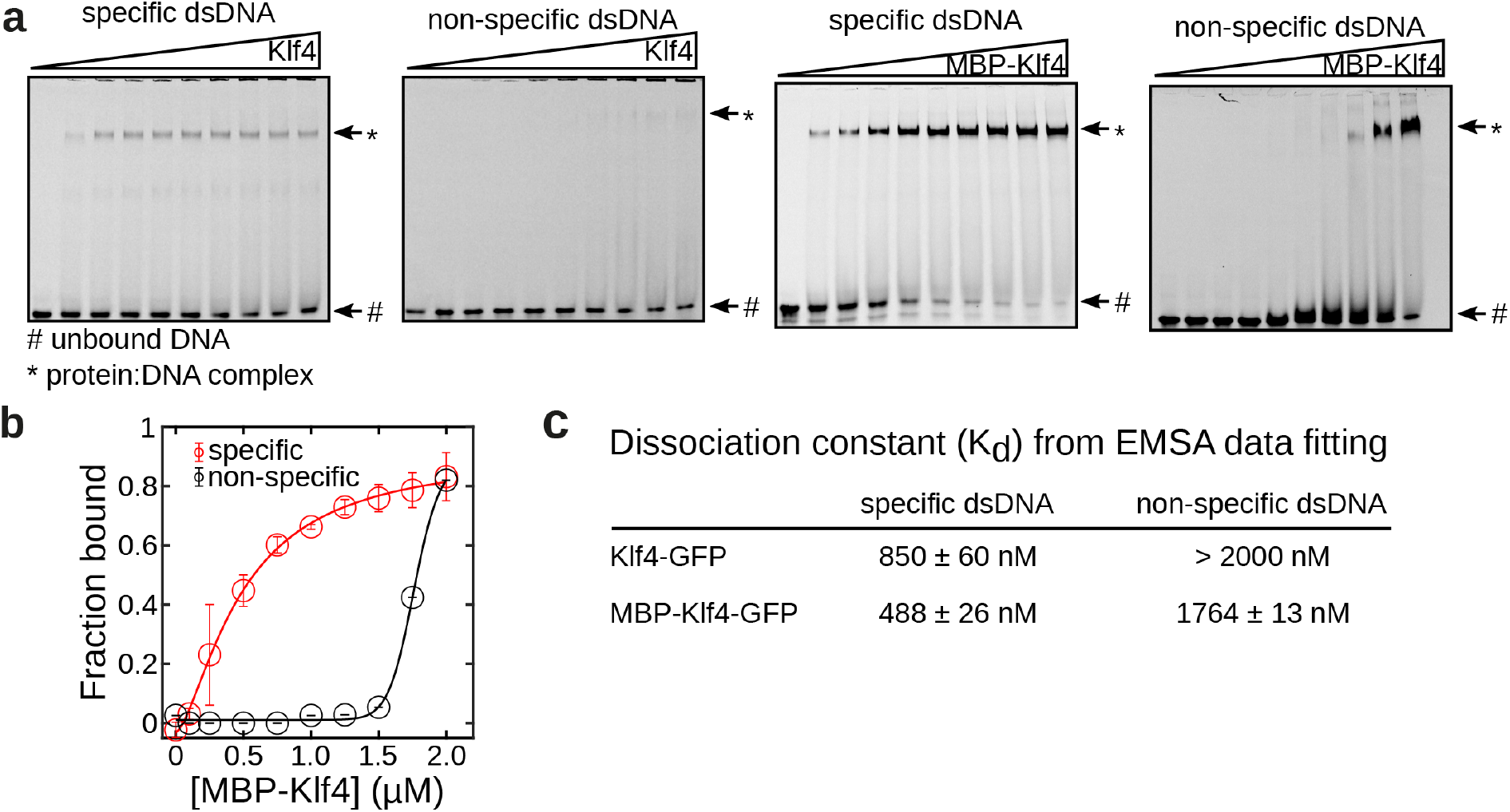
Recombinant pioneer factors bind to DNA in a sequence-specific manner. **a**, Representative examples of TBE gel images visualising Cy5-labelled dsDNA oligonucleotides in electrophoretic mobility shift assays (EMSAs) are depicted. Klf4-GFP and MBP-Klf4-GFP bind to dsDNA in a sequence-specific manner and cause an up-shift of the oligonucleotide in the gel. **b**, EMSAs were performed as shown in **a** to test the affinity of MBP-Klf4-GFP to short dsDNA oligonucleotides with (red) or without (black) specific binding sites for the protein. Quantification of three independent experiments is shown with error bars indicating the standard deviation. **c**, Data fitting of the graphs shown in **b** and Fig. 1b allows determination of the dissolution constant (*K_d_*, see Methods). The error margin indicates the 95% confidence value.

**Extended Data Fig. 3:**
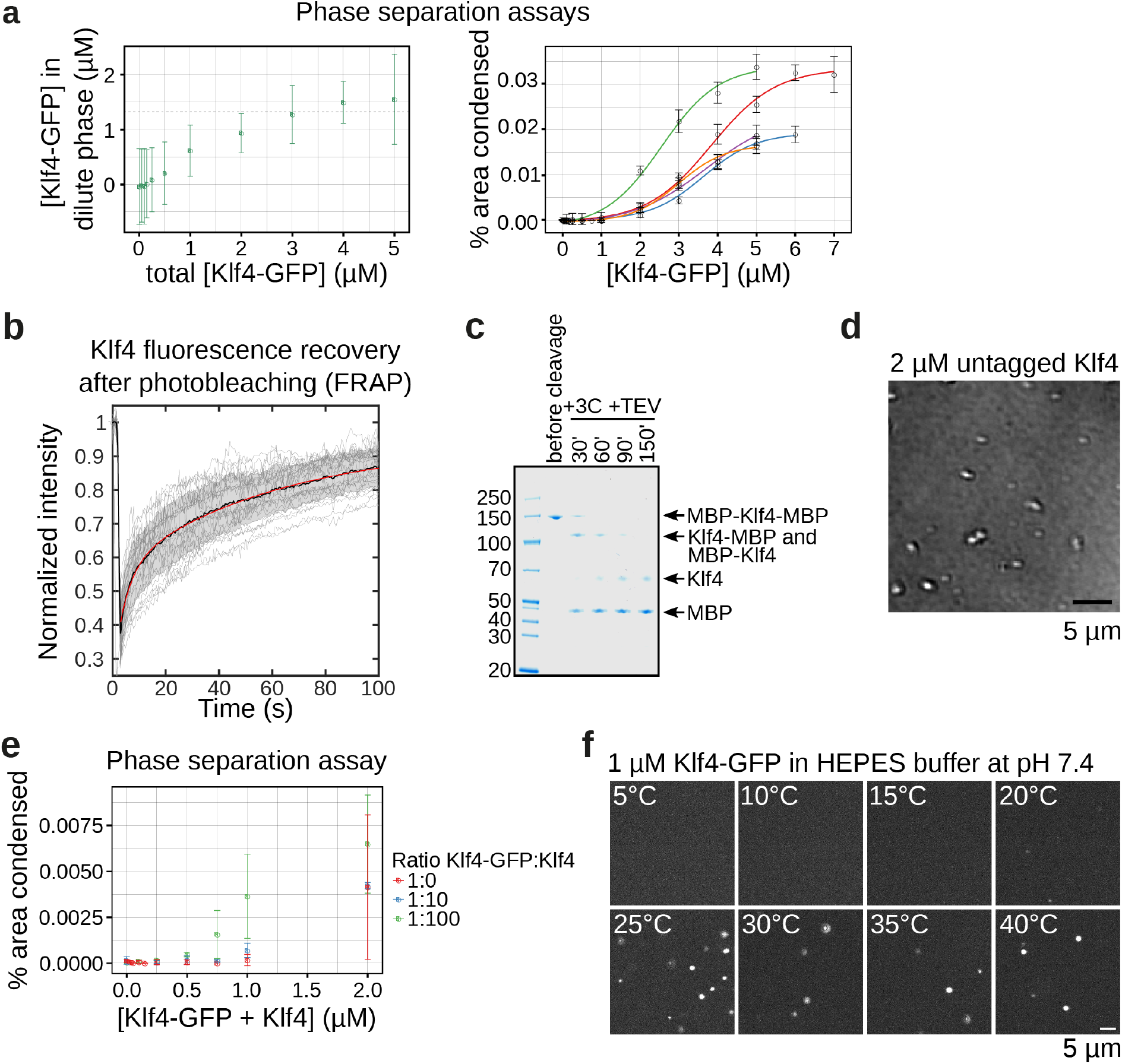
Klf4 forms liquid droplets *in vitro*. **a**, Two kinds of phase separation assays were performed in order to estimate the saturation concentration (*C_SAT_*) for phase separation of Klf4-GFP. Left, the *C_SAT_* is determined by measuring the concentration of the liquid phase after phase separating Klf4-GFP *in vitro* (see Methods). The dashed line indicates the plateau concentration which corresponds to the *C_SAT_* at 1.3 ± 0.3 μM (Mean of 4 repeats; error bars indicate the standard deviation). Right, the fraction of the condensed area in confocal fluorescent microscopy images is quantified. A value above 0 indicates droplets at the given concentration. Here, droplets appear at a concentration above 1 μM. Five repeats are shown as indicated by different colours. **b**, 28 FRAP experiments of Klf4-GFP were performed (grey lines) with the mean depicted as a solid black line (the shaded area sows the 95% confidence value). Data fitting to the mean (red line) was used to estimate a mobile fraction of 93% (see Methods). **c**, In order to test the behaviour of untagged Klf4, MBP-Klf4-MBP was purified with a TEV and 3C PreScission protease cleavage site between the MBP tags and Klf4, respectively. The SDS gel shown depicts a typical purification and cleavage time course and the protein was only used for assays after complete removal of both MBP tags. **d**, Example DIC (differential interference contrast) microscopy image of untagged Klf4 droplets under the same conditions as in Fig. 1c and as used in all other assays. **e**, In order to compare phase separation of untagged Klf4 and Klf4-GFP using confocal fluorescence microscopy, different ratios of Klf4 and Klf4-GFP were phase separated and imaged. The condensed area fraction was used to assess droplet formation as in **a**. For a ratio of 1:0 (100% Klf4-GFP), the mean and standard deviation of the five examples shown in **a** are used. For the 1:10 (10% Klf4-GFP) and 1:100 (1% Klf4-GFP), mean and standard deviation of duplicates are shown. **f**, 1 μM Klf4-GFP was imaged at different temperatures using confocal fluorescence microscopy and a self-built temperature stage which allows rapid temperature changes of a mounted sample. The same field of view is shown at the indicated temperatures. The assay conditions are as for all other assays except that 25 mM HEPES pH 7.4 was used as a buffering component rather than Tris because its pH is less dependent on temperature (△pKa/10°C = −0.14 for HEPES and −0.31 for Tris^77^).

**Extended Data Fig. 4:**
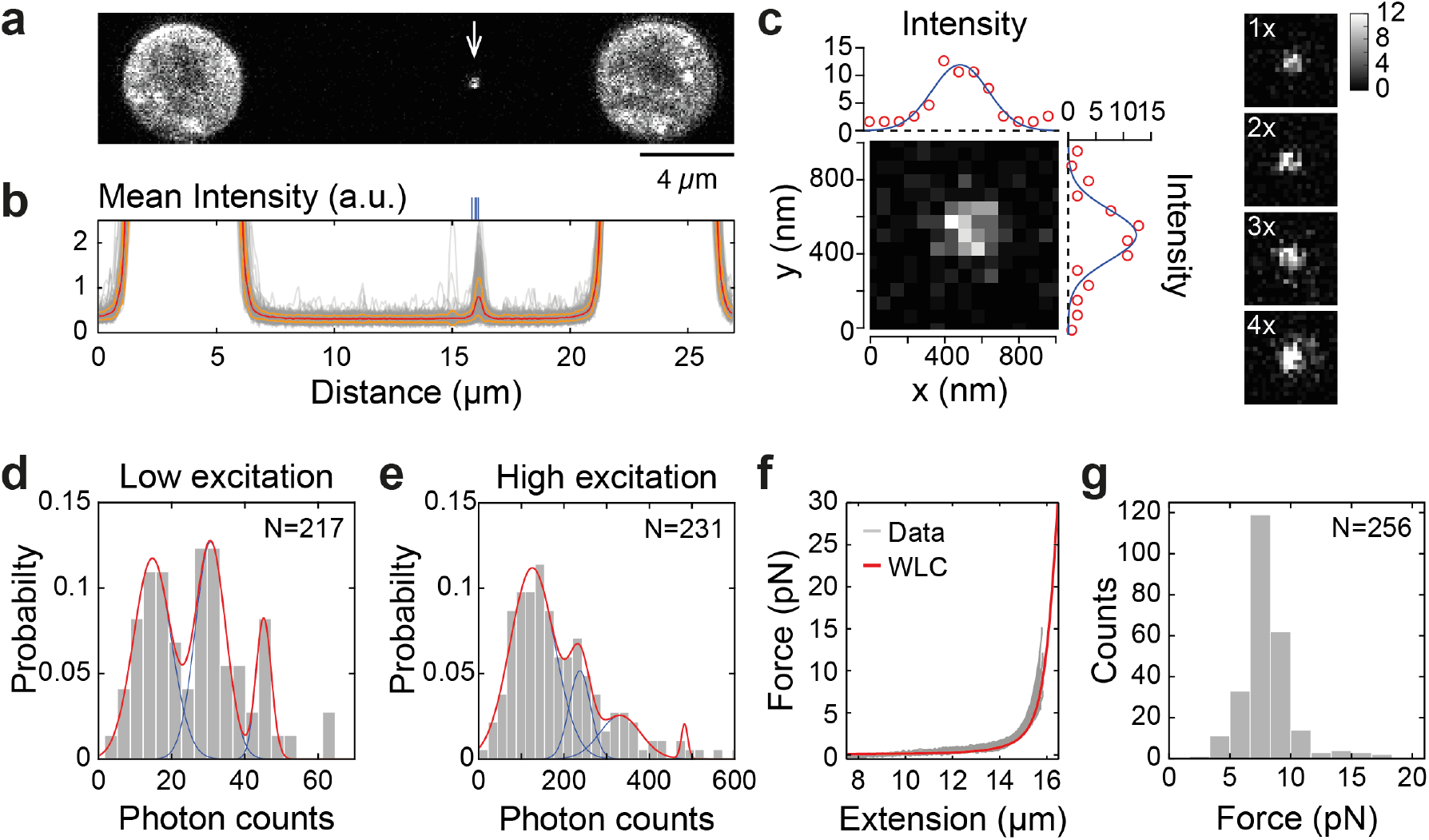
Quantification of the emission intensity per EGFP and determination of the experimental force. **a**, Example confocal image, after background subtraction, of a dCas9-EGFP labelled λ-DNA molecule (the DNA is not labelled) held between two 4.42*μm* diameter polystyrene beads. White arrow shows the position of the dCas9-EGFP probe. **b**, Mean Intensity profile along the DNA. Gray lines correspond to N=270 individual experiments. Red, orange lines correspond to the mean and standard deviation respectively. Blue lines mark the position of the four target sites for dCas9 in λ-DNA (see Extended Data Table 2 and Methods). **c**, Left, example of an individual dCas9-EGFP molecule. The intensity profile in x and y directions (red circles) can be fitted to a gaussian function (solid blue line), rendering a full width at half maximum (FWHM) of 355.2 and 319.5 *nm* in x and y directions respectively. Right, examples of 1, 2, 3 and 4 dCas9-EGFP molecules with integrated intensity values of: 156.1, 237.0, 344.2 and 488.8 photon counts respectively. **d**, **e**, Probability distribution of integrated intensity values (in photon counts units) of dCas9 on λ-DNA in Klf4 assay buffer acquired under low and high intensity imaging conditions respectively (see Methods). The number of experiments in each case (N) is indicated. The distributions were fitted to a gaussian mixture model (3 modes for low and 4 modes for high intensity conditions). Solid red line, sum of the modes, blue lines, individual modes. The emission intensity per EGFP molecule (*I_GFP_*) was determined by taking the average of the differences between the mean intensities of adjacent modes (see Methods and Extended Data Table 4). **f**, A force extension curve (FEC) was recorded prior to each experiment. Gray lines, 30 representative experiments. The experimental FECs are consistent with the theoretical prediction for the force distance curve of a λ-DNA molecule (red line: Worm-like chain model^78^ with contour length of *L_C_* = 16.49 *μm* (48.514 *kbp*), persistence length of *L_P_* = 50 nm and stretch modulus of *K* = 1200 pN). **g**, Histogram of forces at which individual experiments were recorded. *F_exp_* = 8.22 ± 2.65 pN (mean ± standard deviation).

**Extended Data Fig. 5:**
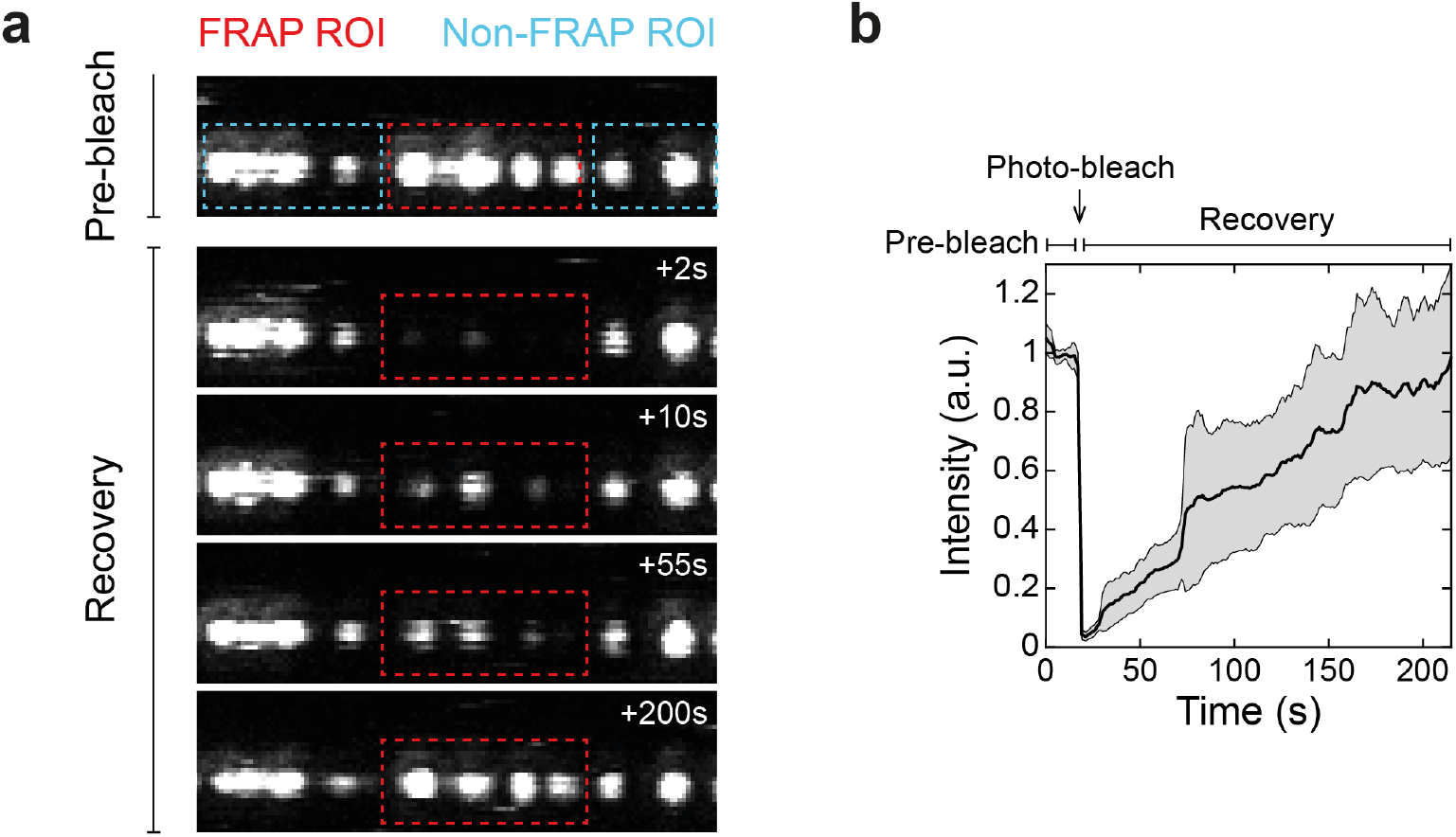
Klf4 fluorescence recovers after photo bleaching. **a**, Representative confocal microscopy images of the time course of a fluorescence recovery after photobleaching (FRAP) experiment. Once the DNA was exposed to a 250 nM concentration Klf4-GFP solution for 200 s (Fig. 2d), the microfluidics chamber was gently flushed. A pre-bleach time series was acquired for 20*s* (last frame shown in top panel), after which, a smaller region of interest (FRAP ROI, red square) was imaged using a high excitation intensity (90% excitation laser power) to induce photo-bleaching. The subsequent recovery of fluorescence was recorded for 200*s*. Time stamps are referenced to the time of induced photo-bleaching. **b**, Mean intensity across the FRAP ROI corrected for photo-bleaching. Individual traces of mean intensity in the FRAP ROI were divided by the mean intensity in the non-FRAP ROI (cyan squares in **a**) and normalized to the value prior to the photo-bleaching step. Black line shows the average trace for N=13 experiments, grey region the standard error of the mean at 95% confidence.

**Extended Data Fig. 6:**
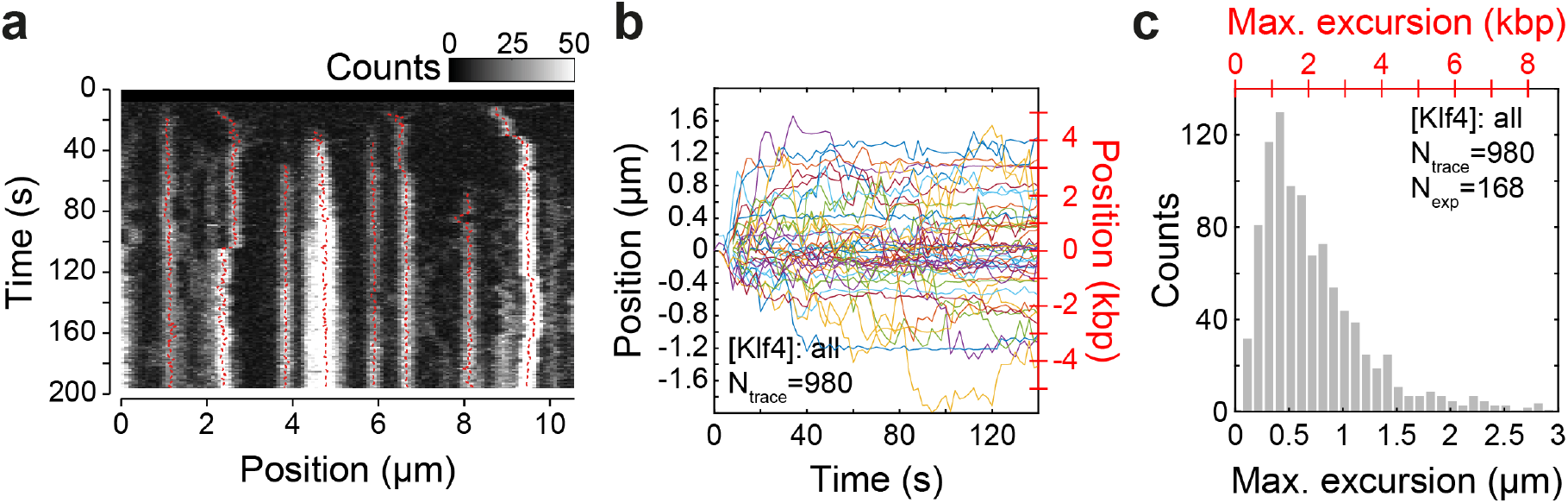
The position of Klf4 clusters fluctuates on DNA. **a**, Kymograph showing the displacement of Klf4-GFP clusters along the DNA. In red, the trajectories of segmented clusters are shown. Segmentation was done using the python package Trackpy (see Methods) with initial search parameters of *P_R_* = 15 pixels, *S_R_* = 6 pixels, *t_memory_* = 30*s* and *t_min_* = 100*s*. **b**, Position of foci over time along the DNA. Position is plotted relative to the initial point of each trace (*Position* = *x*(*t*) – *x*(*t* = 0), where *x*(*t*) is the position over time). A random selection of 50 traces is shown. **c**, Histogram of the maximum excursion length (defined as the farthest point a cluster moves from the initial position (*Max. excursion* = *max*(|*x*(*t*) – *x*(*t* = 0)|))). The number of traces (*N_trace_*) is indicated. *N_exp_* = 168 experiments were considered in the concentration range: [*Klf*4] = 3 – 281n*M*.

**Extended Data Fig. 7:**
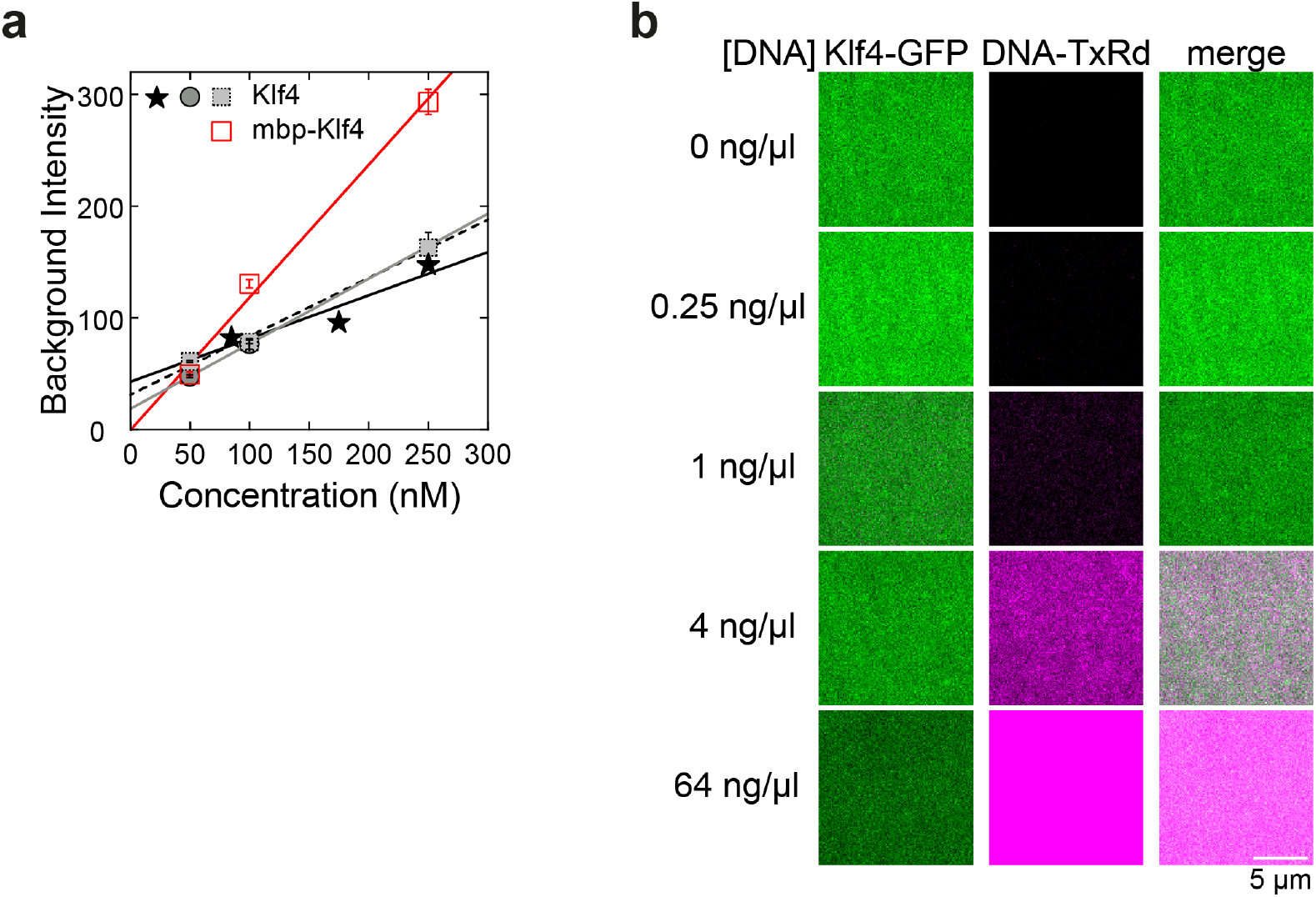
Klf4 concentration remains below *C_SAT_* in the optical tweezers assay and Klf4-GFP droplets are not formed via a nucleation mechanism. **a**, Background intensity, computed as the mean intensity in an ROI distant from the DNA, as a function of the loaded concentration. Each data set corresponds to a dilution series starting from the highest concentration, following the indicated concentration points. Symbols represent the mean of the last 3 experiment of each series. Error bars show the standard deviation. A linear fit rendered the slope (*I_nM_* in units of Intensity / nM) and the intercept (*I*_0_ in intensity units). See results in Extended Data Table 3. **b**, Confocal fluorescence images depict a phase separation assay with 500 nM Klf4-GFP and increasing amounts of Sulforhodamine 101-X (TxRd)-labelled dsDNA oligonucleotides that contained 9 binding sites for Klf4. In contrast to lambda-DNA (Fig. 1c), the short oligonucleotides fail to induce Klf4 condensation, indicating that Klf4 condensates do not form on DNA via a mechanism in which DNA serves as a nucleator for phase separation. All confocal microscopy images of the same color have the same contrast settings.

**Extended Data Fig. 8:**
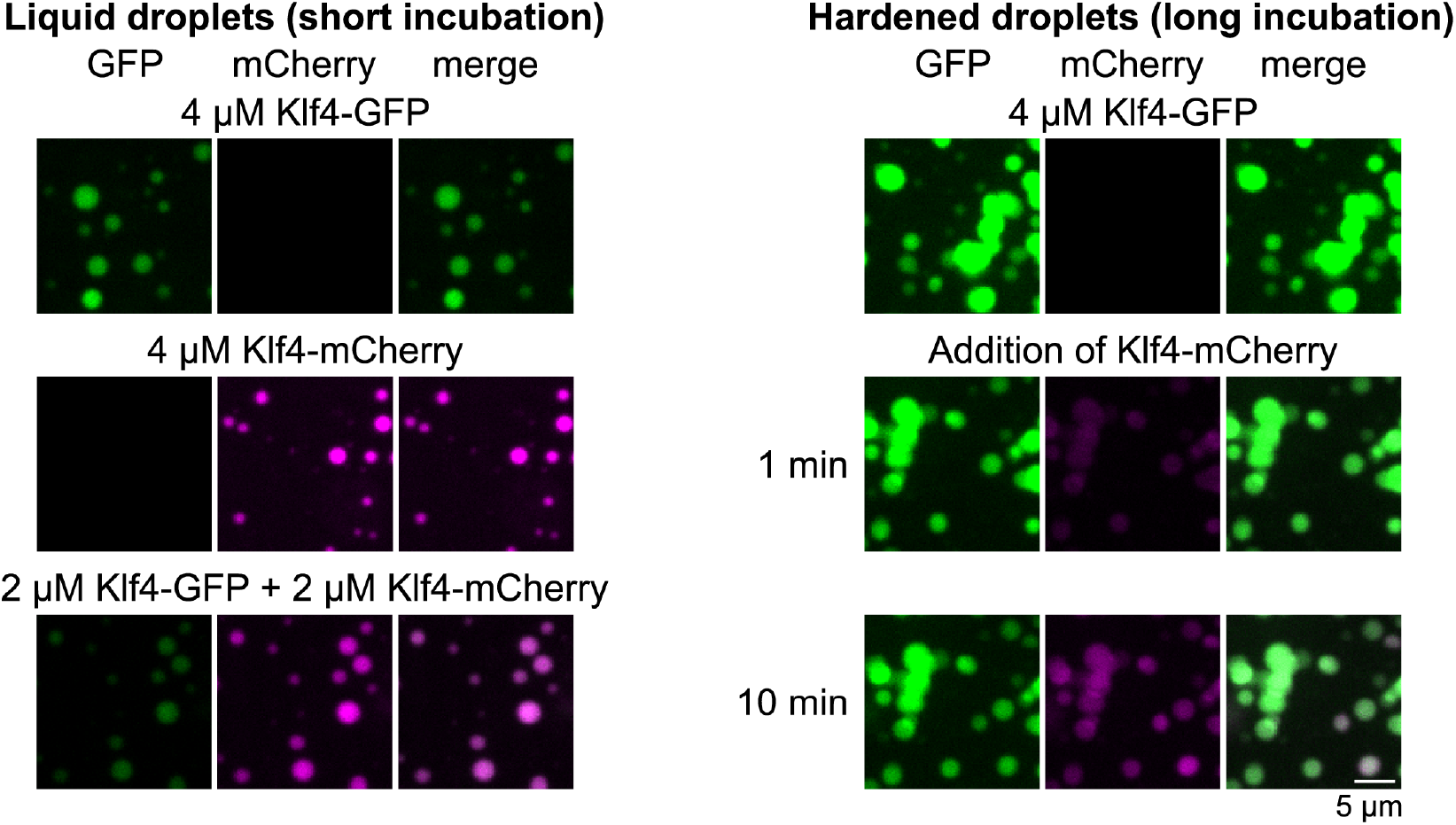
Hardened Klf4 droplets are able to grow. In order to exclude the possibility that Klf4 droplets stopped growing on DNA because of hardening, we tested whether hardened droplets retain the ability to grow. Right, for this, Klf4-GFP droplets were formed and incubated until hardened, which is evident from the fact that the droplets stopped fusing and are instead sticking together (top row). After 50 min, a fresh solution of Klf4-mCherry was added (middle row). After only 10 min (bottom row), the green Klf4-GFP droplets enriched red signal of Klf4-mCherry. Left, as a control, to test whether both Klf4-GFP and Klf4-mCherry individually (top and middle row) or in combination (bottom row) are able to form droplets under these conditions, the standard incubation time for droplet assays was used (short incubation: 20 min). See Methods for details. All confocal microscopy images of the same color have the same contrast settings.

**Extended Data Fig. 9:**
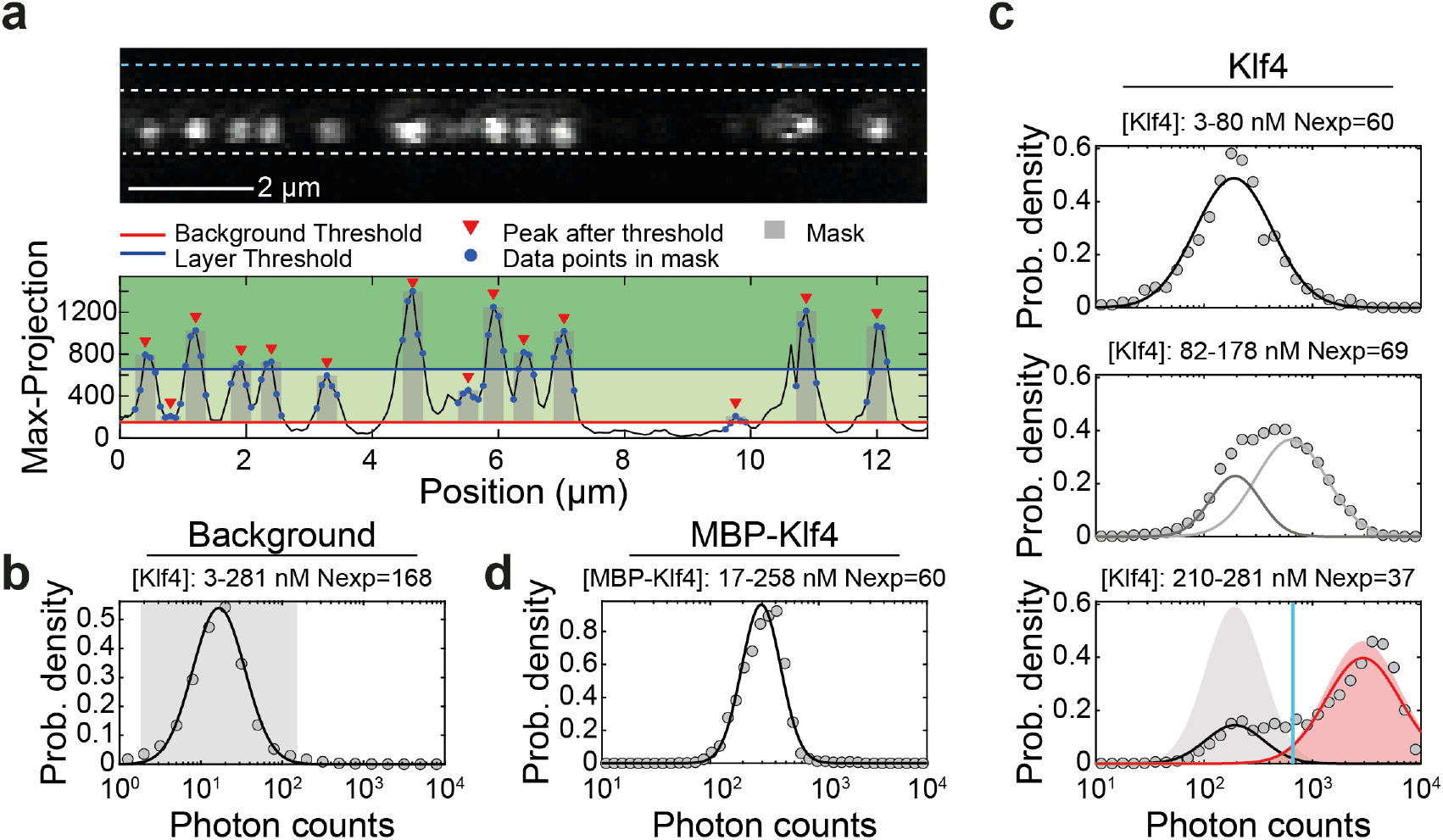
Determination of the intensity thresholds for pixel classification into adsorbed and condensed states. **a**, Top, example confocal image of Klf4-GFP on DNA. Bottom, intensity profile and data processing steps for the experiment shown in top. First, we extracted the maximum projection intensity profile along the DNA (black curve). The profile is determined in a region of 20 pixels around the DNA axis (white dashed lines in top). We next subtracted the background (see below) and filterd the data with a spacial mask: we applied the findpeaks algorithm from Matlab (peak positions marked with red triangles) and selected data points between –*dr* and +*dr* from the center of the peaks with the only restriction that two intervals cannot overlap (blue points and gray area show the result for *dr* = 2). Data points selected this way were used in the determination of the intensity distributions shown in Fig. 3b. **b**, We extracted the background values (after background subtraction) along a line away from the DNA (cyan dashed line in **a**, top). The probability density of the logarithm of pixel intensities, pulling together all Klf4-GFP experiments, can be fitted to a normal probability density function (black line). The background threshold (*I_th–bg_* = 152.9, red line in **a**, bottom) was defined as the mean plus 3 times the standard deviation of the distribution (gray area, upper boundary). **c**, Probability density of the logarithm of pixel intensities, along the intensity profiles pulling together Klf4-GFP experiments in the concentration range indicated. For low protein concentrations (top), the probability density can be fitted to a normal probability density function (black line, mean *μ*_1_ = 187.96 and standard deviation *σ*_1_ = 237.47). For high protein concentrations (bottom), the probability density was fitted to a two components gaussian mixture model constraining the mean of the first mode to the value extracted from the low concentration distribution in the top pannel (black line *σ*_1_ = 162.50, area *a*_1_ = 0.22 and red line, *μ*_2_ = 2.93 * 10^3^, *σ*_2_ = 3.52 * 10^3^, *a*_2_ = 0.78 respectively). We define the adsorption layer upper boundary (*I_th–ads_* = 658.5, cyan line) as the crossing point between the first and second modes, normalized to the same area independently (gray and red areas, rescaled here for representation purposes: maximum value = 0.65 and 0.50 respectively). In the intermediate concentration range, an unconstrained fit to a two components gaussian mixture model rendered a low intensity component with a mean similar to the one observed at low and high concentrations (dark gray line, *μ*_1_ = 193.63, *σ*_1_ = 136.37, *a*_1_ = 0.31) and a high intensity component (light gray line, *μ*_2_ = 634.66 and *σ*_2_ = 708.06, *a*_2_ = 0.69)). **d**, The probability density pulling together all MBP-Klf4 experiments, can be fitted to a normal probability density function rendering a mean value similar to the one found for the adsorption state of Klf4-GFP (black line, *μ* = 295.40 and *σ* = 151.74).

**Extended Data Fig. 10:**
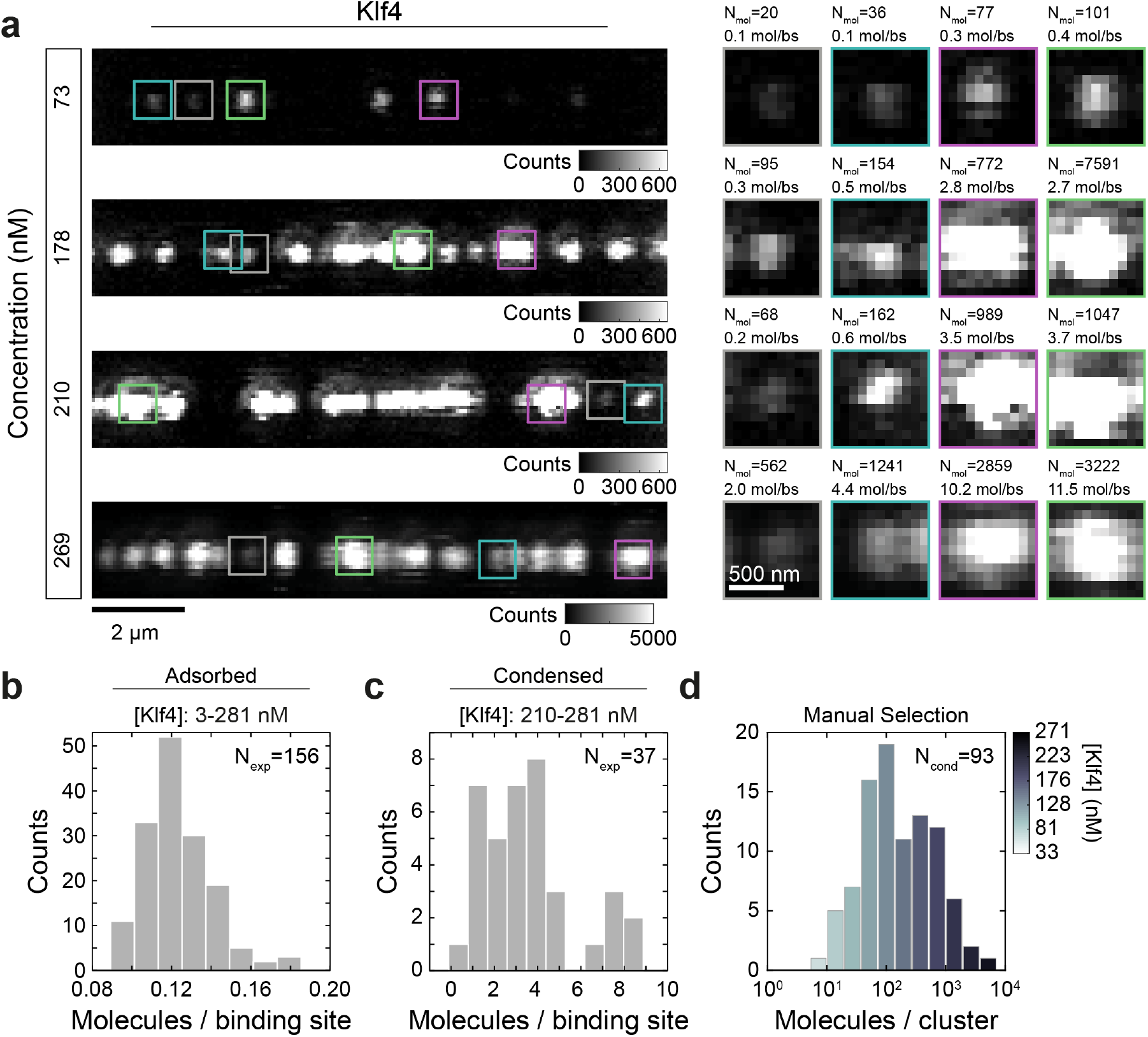
Number of molecules per binding site in the adsorbed and condensed states for Klf4. **a**, Left: Representative confocal images of Klf4-GFP coated DNA at the specified bulk concentration. Right: For each experiment, a selection of four clusters is shown, with the corresponding number of molecules per cluster and the resulting number of molecules per binding site. The number of molecules per cluster (*N_mol_*) was determined as the integrated intensity across the ROI divided by the intensity per GFP. The number of molecules per binding site was computed as the number of molecules divided by the number of biding sites (of 10 base pairs length) for the considered length (see Methods). The top three rows are displayed in the same intensity scale, which is set to a maximum value coincident with the upper intensity threshold of the adsorption state (*I_ads–max_* = 658.5). **b, c**, Number of molecules per binding site for the adsorbed and condensed regions. Here, the number of molecules that correspond to the adsorbed (condensed) state was determined as the sum of the integrated intensity corresponding to the number of pixels classified as adsorbed (condensed) divided by the intensity per GFP. Finally, the number of molecules per binding site is obtained by taking into account the length of a binding site. The corresponding concentration range and number of experiments (*N_exp_*) is indicated. **d**, Histogram of number of molecules per cluster at different Klf4 concentrations. For each experiment, isolated clusters were manually selected when possible. The number of molecules (*N_mol_*) was determined as in **a**.

**Extended Data Fig. 11:**
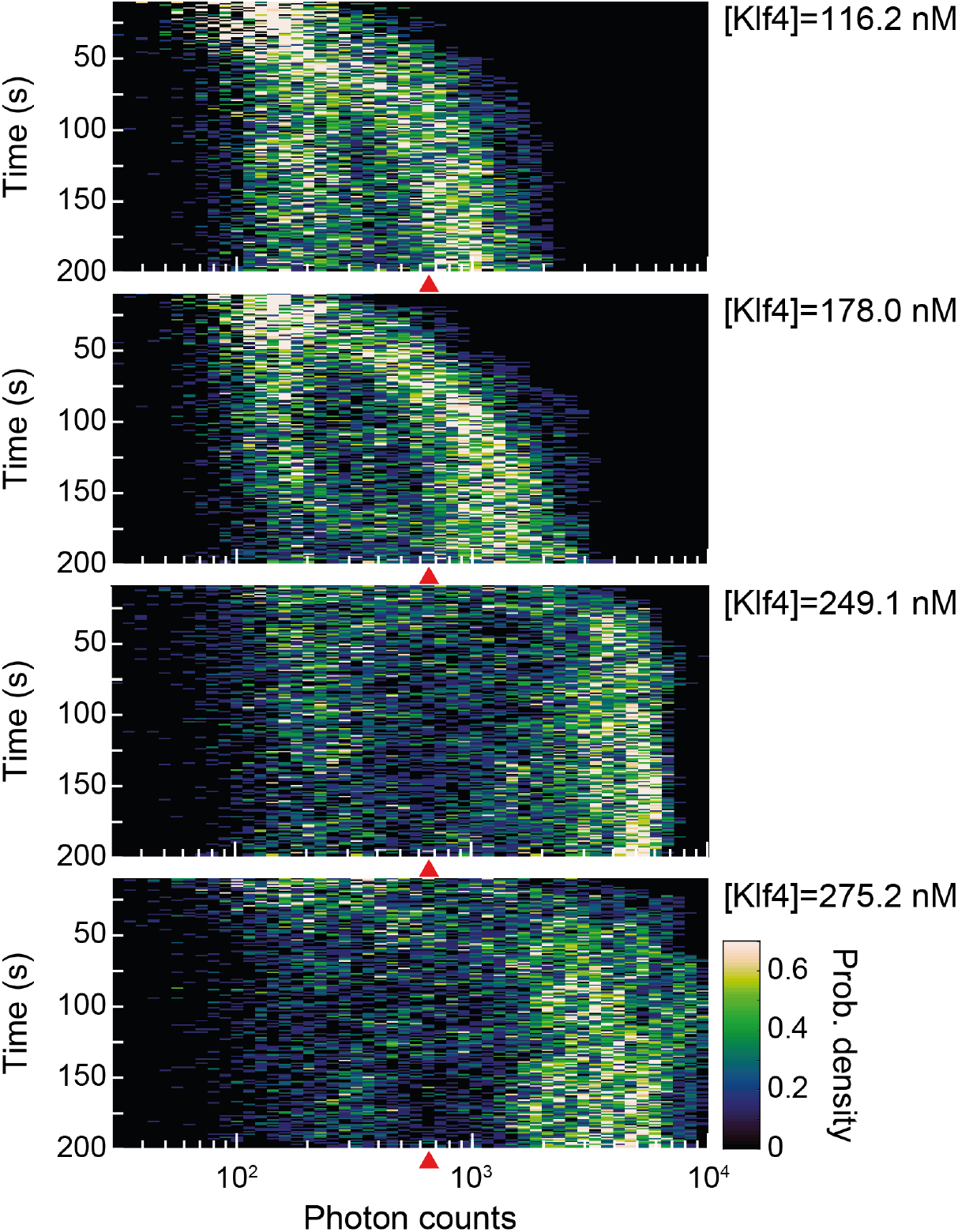
Representative examples of the time evolution of the intensity distributions for Klf4. Probability density of the logarithm of intensities as a function of time after exposure of DNA to Klf4-GFP for individual experiments recorded at the specified Klf4-GFP bulk concentration. Intensity distributions were computed for each frame as described in Extended Data Fig. 9. Red triangle, position of the adsorption threshold used in pixel classification (Fig.3b and Extended Data Fig. 9). After an initial binding step (first ~ 50s in top) the intensity distribution bifurcates into the adsorbed and condensed states (low and high intensity branches respectively). Over time, these states coexist and the condensed one increases its brightness (the high intensity branch migrates toward higher intensity values) until it reaches a stable value (~ 10^3^ photon counts in top). Remarkably, the way these states are populated are suggestive of a switch like behavior; the condensed state becomes more populated over time at the expense of the adsorbed one. Increasing the bulk concentration reduces the time required to observed the bifurcation. Moreover at higher bulk concentrations the condensed state gets brighter (~ 5 * 10^3^ in bottom) and overall more populated.

**Extended Data Fig. 12:**
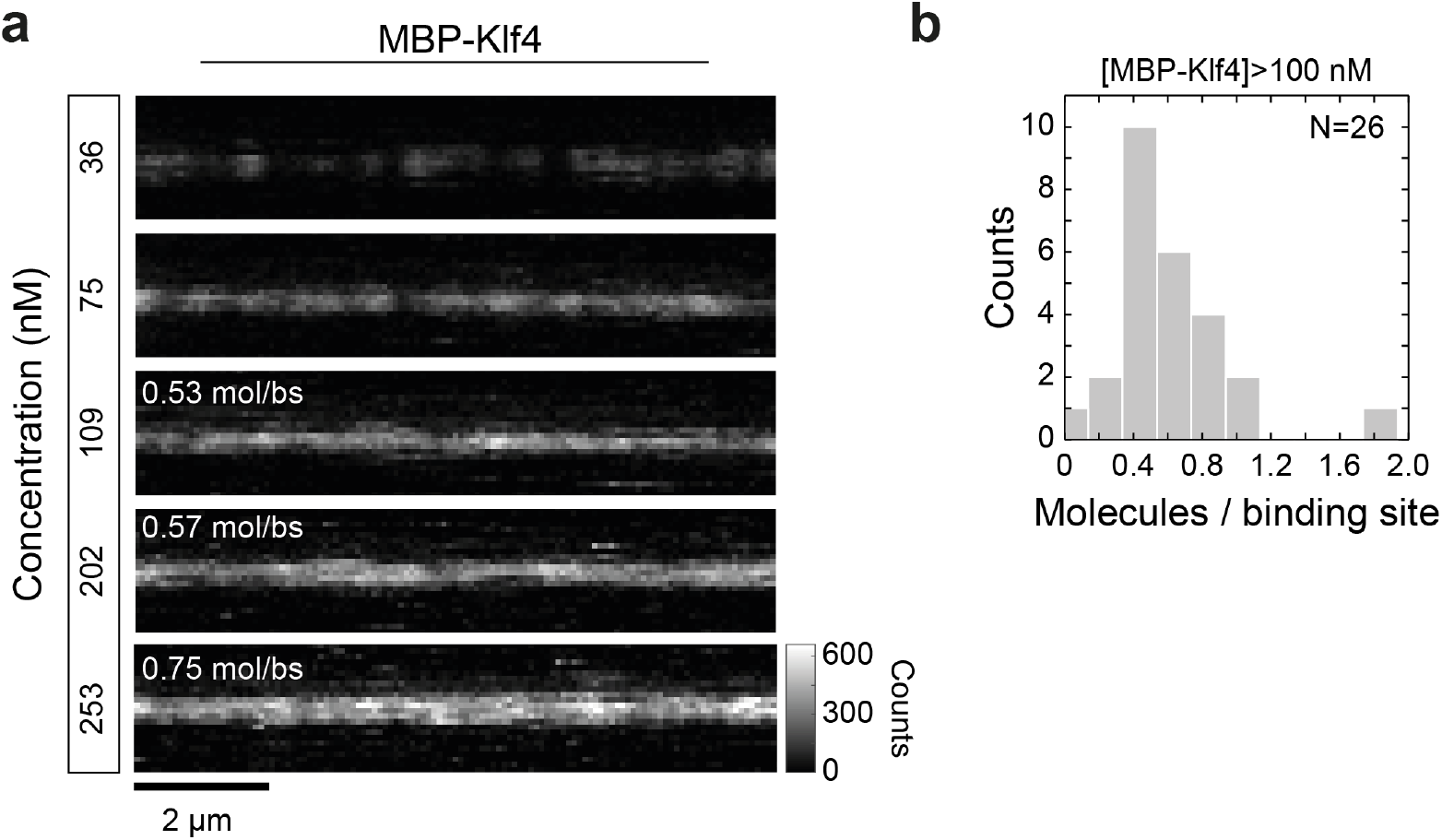
Number of molecules per binding site for MBP-Klf4. **a**, Representative confocal images of MBP-Klf4-GFP coated DNA at the specified bulk concentration. The images are displayed in the same intensity scale, which is set to a maximum value coincident with the upper intensity threshold of the adsorption state for Klf4 (*I_ads–max_* = 658.5). **b**, Number of molecules per binding site computed as in Extended Data Fig. 10. Only experiments that exhibit full coverage (all experiments for [*MBP – Klf*4] > 100nM, N = 26) were considered for this calculation.

**Extended Data Fig. 13:**
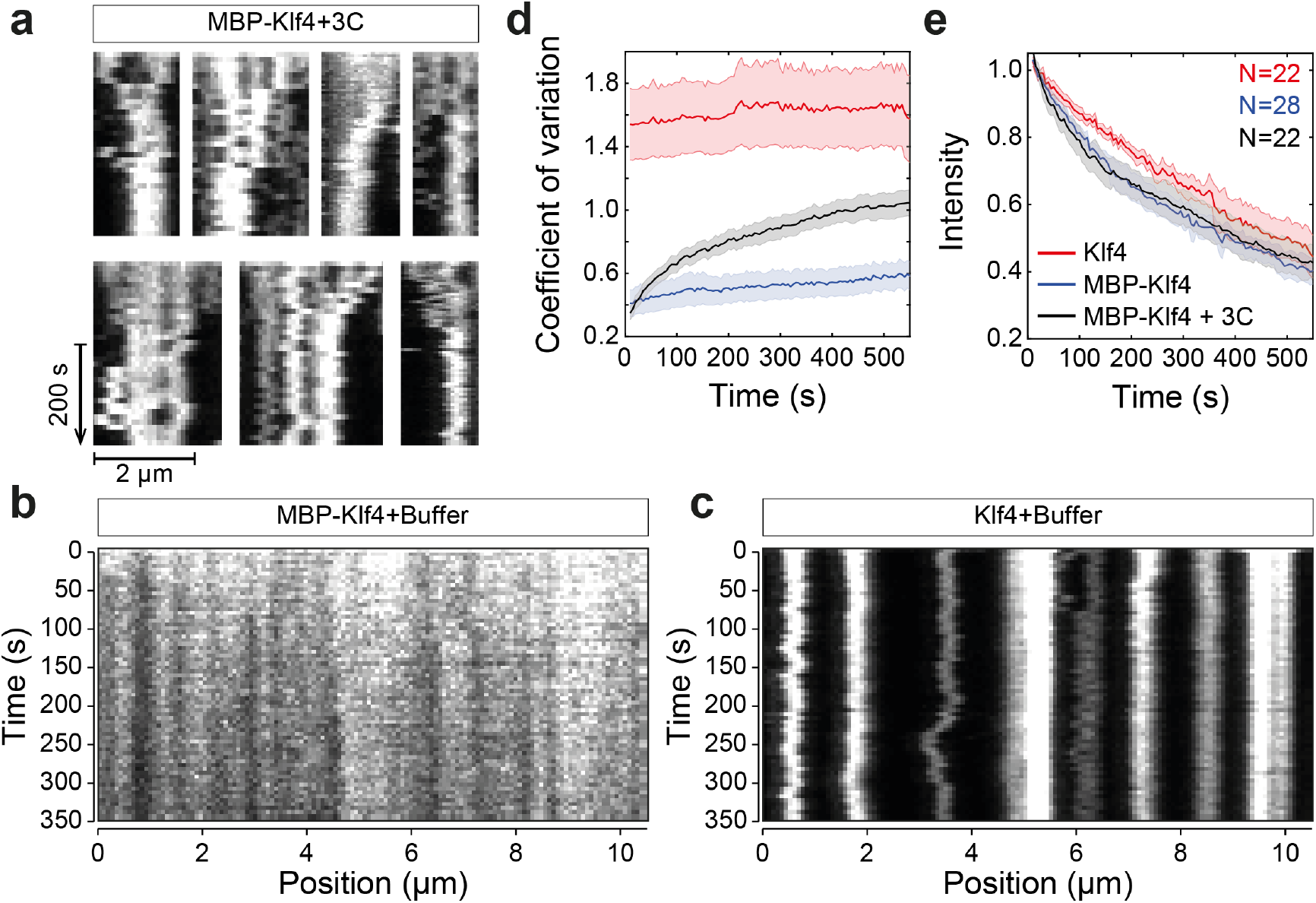
Klf4 transition from adsorbed to condensed layer on the DNA upon MBP cleavage. **a**, Representative examples of kymograph ROIs corresponding to the 3C dependent MBP-Klf4 condensation. After the DNA chain is coated with MBP-Klf4, the DNA-beads system is transferred to a solution containing the 3C protease and no free MBP-Klf4 in solution (Fig. 3f). Over time, the thin layer coating the DNA undergoes a protease dependent rearrangement on the chain to form several condensed spots (local increase in intensity). Each example corresponds to a different experimental realization. **b, c**, Example kymographs of MBP-Klf4 and Klf4 coated DNA when transferred to the assay buffer (which does not contain the 3C protease). In this case, the spatial arrangement of intensities along the DNA remains unchanged **d**, The coefficient of variation (CV) of the distribution of intensity values along the DNA is shown. For the cases in which the system is transferred to a protease free buffer, the CV remains constant over time, while in the presence of the protease it increases. **e**, Notably, the average intensity on the chain decays in a similar way for the three situations. This result is consistent with a rearrangement of material on the DNA that leads to local enrichment (as well as local depletion) of cleaved Klf4-GFP. Solid lines, mean and shaded areas the standard error of the mean at 95% confidence. The number of experiments in each case is indicated.

**Extended Data Fig. 14:**
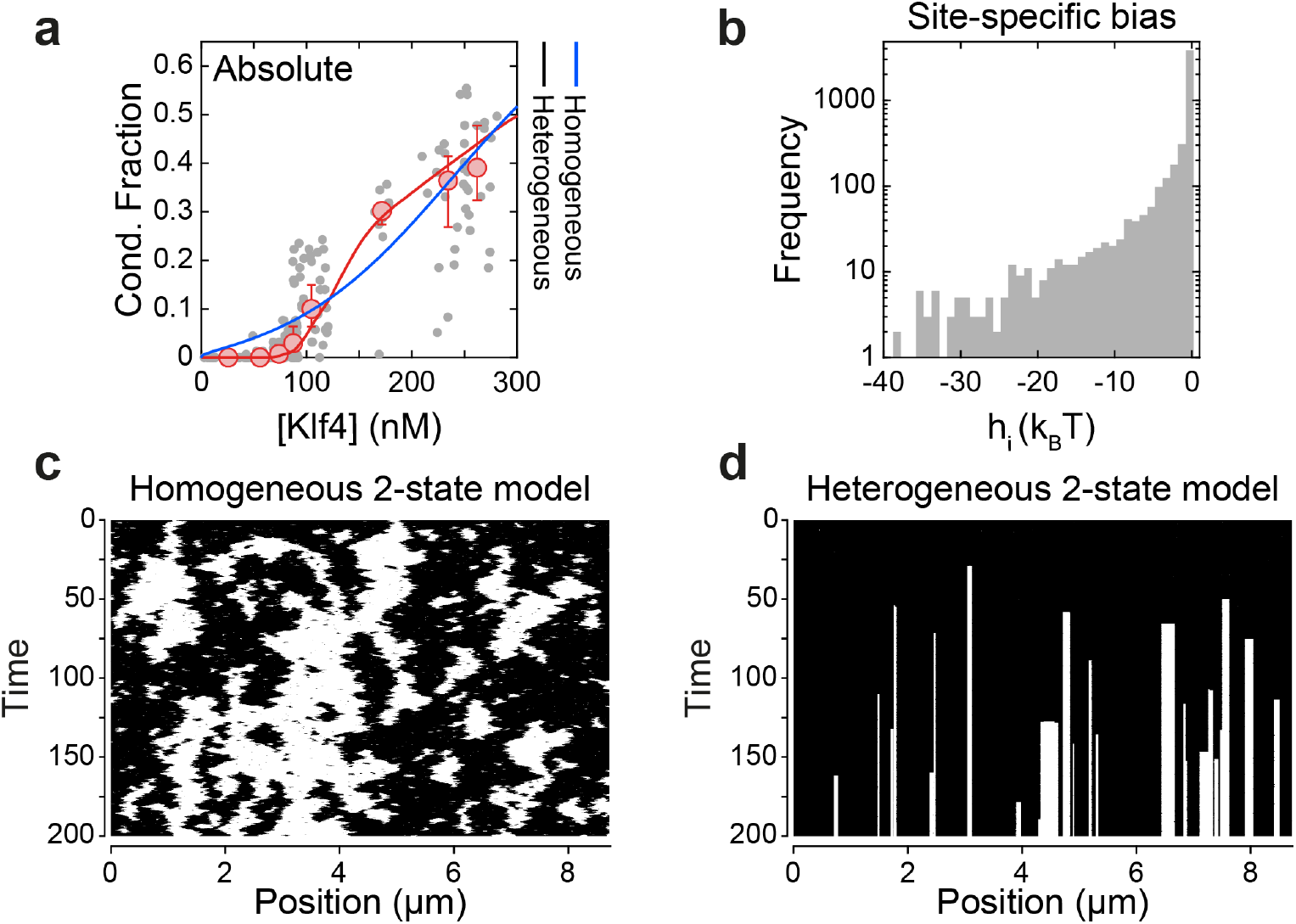
A heterogeneous two state model can successfully account for experimental observations. **a**, Data for the condensed fraction for Klf4-GFP at different concentrations are shown (same as in Fig. 3d). Blue and red lines are fits to the homogeneous and the heterogeneous models respectively. From the fit to the homogeneous model, we extract the following parameter values: *J* = 3.37 *kT, C*_0_ = 292 *nM*, and *α* = 0.0016 *kT*. For the heterogeneous model, the corresponding parameter values are *J* = 6 *kT, C*_0_ = 35 *nM, α* = 0.5 *kT, k* = 0.1, and *θ* = 18 kT (see Methods). **b**, Histogram of *h_i_* i.e. the site-specific propensity for condensation is shown. This histogram is gamma distributed with shape and scale parameters *k* = 0.1, and *θ* = 18 *kT* respectively. **c**, A representative kymograph is shown for the homogeneous model using the fitted parameter values, corresponding to a concentration of 200 nM. It was generated by simulating the kinetics of the model using Gillespie simulations (see Methods). Evidently, the model fails to capture the length scales of condensed and adsorbed layers. **d**, Example kymograph for the heterogeneous model is shown for a concentration of 200 *nM*. This model can capture the dynamics of the condensed layers as observed in experiments.

## Extended Data Tables 1-4

**Extended Data Table 1:**
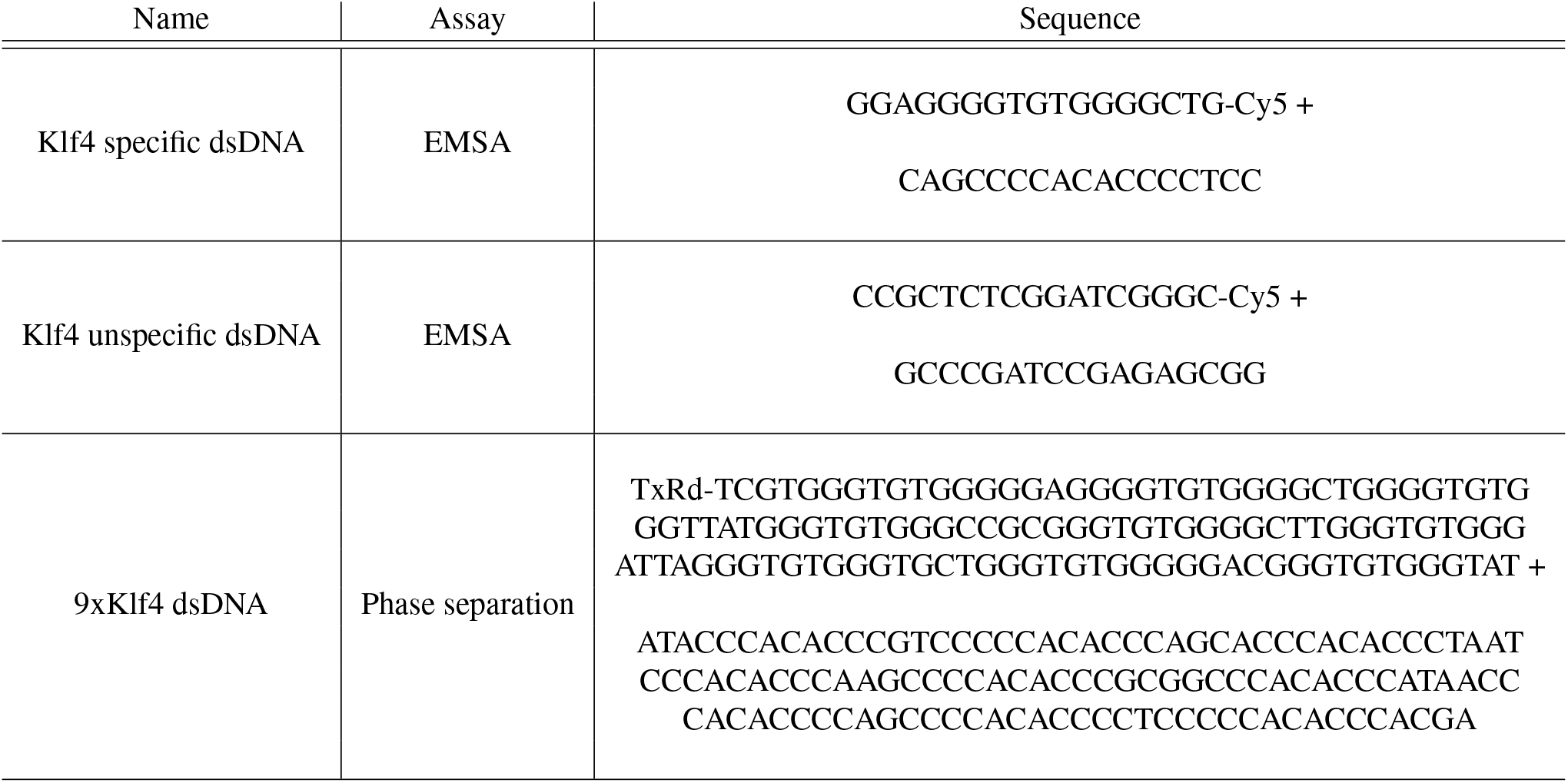
Oligos used in this study.

**Extended Data Table 2:**
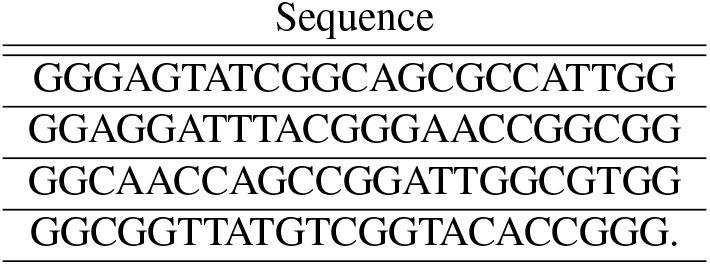
Target sequences for dCas9 on λ-DNA.

**Extended Data Table 3:**
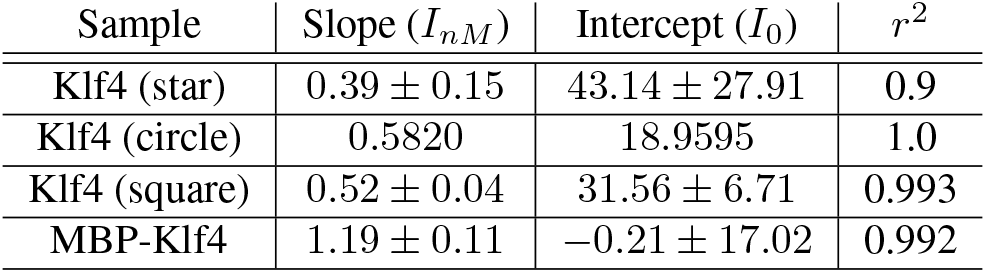
Linear fit to the background intensity vs concentration curve for all dilution series. Standard error of the parameters is indicated.

**Extended Data Table 4:**
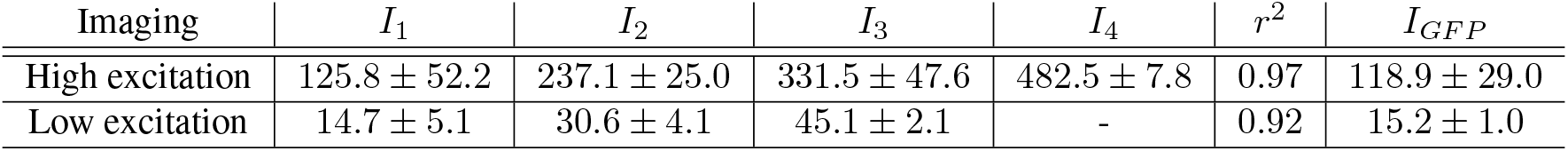
Emission intensity of individual EGFP molecules. Mean ± standard deviation extracted from a fit of the integrated intensity values of dCas9-EGFP to a gaussian mixture model.

| Imaging | $I_1$ | $I_2$ | $I_3$ | $I_4$ | $r^2$ | $I_{GFP}$ |
| --- | --- | --- | --- | --- | --- | --- |
| High excitation | $125.8 \pm 52.2$ | $237.1 \pm 25.0$ | $331.5 \pm 47.6$ | $482.5 \pm 7.8$ | 0.97 | $118.9 \pm 29.0$ |
| Low excitation | $14.7 \pm 5.1$ | $30.6 \pm 4.1$ | $45.1 \pm 2.1$ | - | 0.92 | $15.2 \pm 1.0$ |

